# Genome-wide analysis of long terminal repeat retrotransposons from the cranberry *Vaccinium macrocarpon*

**DOI:** 10.1101/2021.07.15.452536

**Authors:** Nusrat Sultana, Gerhard Menzel, Kathrin M. Seibt, Sònia Garcia, Beatrice Weber, Sedat Serçe, Tony Heitkam

## Abstract

**BACKGROUND:** Long terminal repeat (LTR) retrotransposons are widespread in plant genomes and play a large role in the generation of genomic variation. Despite this, their identification and characterization remains challenging, especially for non-model genomes. Hence, LTR retrotransposons remain undercharacterized in *Vaccinium* genomes, although they may be beneficial for current berry breeding efforts.

**OBJECTIVE:** Exemplarily focusing on the genome of American cranberry (*Vaccinium macrocarpon* Aiton), we aim to generate an overview of the LTR retrotransposon landscape, highlighting the abundance, transcriptional activity, sequence, and structure of the major retrotransposon lineages.

**METHODS:** Graph-based clustering of whole genome shotgun Illumina reads was performed to identify the most abundant LTR retrotransposons and to reconstruct representative *in silico* full-length elements. To generate insights into the LTR retrotransposon diversity in *V. macrocarpon*, we also queried the genome assembly for presence of reverse transcriptases (RTs), the key domain of LTR retrotransposons. Using transcriptomic data, transcriptional activity of retrotransposons corresponding to the consensuses was analyzed.

**RESULTS:** We provide an in-depth characterization of the LTR retrotransposon landscape in the *V. macrocarpon* genome. Based on 475 RTs harvested from the genome assembly, we detect a high retrotransposon variety, with all major lineages present. To better understand their structural hallmarks, we reconstructed 26 Ty1-*copia* and 28 Ty3-*gypsy in silico* consensuses that capture the detected diversity. Accordingly, we frequently identify association with tandemly repeated motifs, extra open reading frames, and specialized, lineage-typical domains. Based on the overall high genomic abundance and transcriptional activity, we suggest that retrotransposons of the Ale and Athila lineages are most promising to monitor retrotransposon-derived polymorphisms across accessions.

**CONCLUSIONS:** We conclude that LTR retrotransposons are major components of the *V. macrocarpon* genome. The representative consensuses provide an entry point for further *Vaccinium* genome analyses and may be applied to derive molecular markers for enhancing cranberry selection and breeding.

## INTRODUCTION

The genus *Vaccinium* L. belongs to the family Ericaceae Juss. comprising approx. 450 species distributed all over the world [1]. Available molecular phylogenies suggest that the genus is monophyletic, whereas intrageneric relationships are more difficult to resolve [2, 3]. Dependent on different systematic treatments, the genus is divided into approximately 30 sections and subgenera [1–4]. *Vaccinium macrocarpon* Aiton, also known as the American cranberry, is native to North America [5, 6] and is one of the most commonly cultivated *Vaccinium* species, along with the highbush blueberry (*V. corymbosum* L.) [5]. The small berry fruits produced from wild and cultivated species of the genus are rich in nutritious secondary plant metabolites, including some beneficial for human health with anticancer, antioxidant, antidiabetic, and many other properties [7, 8]. The American cranberry and the European cranberry (a close relative, *V. oxycoccus* L.) belong to the same section or subgenus, named *Oxycoccus* [1, 4]. Although these two species differ in geographical distribution, they can produce fertile offspring upon hybridization [5, 6].

*Vaccinium* species have ploidy levels ranging from diploid to hexaploid, with a base chromosome number x = 12 [5]. As cranberry (*V. macrocarpon*) is a diploid and self-fertile species with a relatively small genome size (approximately 470 Mb), it has become a valuable reference to study the genetics and genomics of the genus [6, 9–11]. So far, whole genome sequence data, a reference genome assembly, and transcriptome datasets have been published for the diploid cranberry and for both the diploid and the tetraploid blueberries [11–13]. In addition, several high-throughput genetics, genomics, and transcriptomics datasets as well as other important breeding information of different species of the genus *Vaccinium* are either published or underway, accessible through the web platform https://www.vaccinium.org.

Recent technological advancements allow tracing of genome dynamics on a fine-scale level. Especially transposable elements (TEs) play a major role in genome evolution and contribute largely to the generation of polymorphisms between genotypes [14–18]. Hence, TEs can also be applied as molecular markers for the differentiation of accessions and to inform plant breeding programs [19–23]. Thus, the correct annotation and characterization of repetitive sequences, particularly TEs, are a prerequisite for an enriched genome assembly as well as for subsequent genome characterization and valorization [10–12]. Nevertheless, as many different TE classification hierarchies have been suggested [24–29], as an integration of tools, databases, and pipelines is still discussed [30], and as there is still no gold standard for TE identification and classification available, TE annotation and classification for non-model species is still a considerable undertaking.

In plants, the most abundant TEs belong to the group of long terminal repeat (LTR) retrotransposons contributing up to 80 % of nuclear DNA [31–33]. They consist of two LTRs flanking the main protein-encoding genes *gag* and *pol* [25, 33, 34]. Whereas *gag* (encoding the capsid and nucleocapsid) is considered as a single gene, the *pol* gene generally encodes several domains, including a protease (PR), an RNase H (RNH), a reverse transcriptase (RT), and an integrase (INT) [25, 33, 34]. In addition to the protein domains, two conserved motifs are characteristic for these full-length elements: a primer binding site (PBS) and a polypurine tract (PPT) [25, 31, 33, 34]. Based on the order of the protein domains and their sequence, the LTR retrotransposons are further divided into the superfamilies Ty3-*gypsy* and Ty1-*copia* [24, 28].

In *Vaccinium* genomes, we have already gained first insights into the repetitive DNA composition, especially into that of satellite DNAs [35, 36]. For dispersed repeats, first estimates were also brought forward, suggesting that nearly 40 % of the cranberry genome was composed of TEs [10]. Very recently, a TE annotation was also made available for the genome of tetraploid blueberry (*V. corymbosum*) [13]. However, the contribution of individual TE families to these genomes is still not well-studied [10–12].

Here, we aim to complement the broad overview studies with a deep analysis of the most abundant, individual TE families in the cranberry genome. The precise knowledge of the LTR retrotransposon composition, as well as the underlying sequences and structures can inform annotation of berry genomes and may provide support for berry breeding and selection.

## MATERIALS AND METHODS

### Graph-based clustering of repeat sequences of the *V. macrocarpon* genome

To representatively survey the cranberry LTR retrotransposon landscape, we used publicly available whole genome paired-end Illumina reads of *Vaccinium macrocarpon* (NCBI BioProject PRJNA245813) of cranberry cultivar ‘Ben Lear’ (accession number CNJ99-125-1) [10]. Before clustering, we pre-treated the reads by quality filtering, adapter trimming, and processing of read length: Illumina Truseq adapters were removed using Trimmomatic with the parameters ILLUMINACLIP:TruSeq3-PE-2.fa:2:30:10 [37]. Fastx_trimmer and FASTx quality filter from FASTX-Toolkit (http://hannonlab.cshl.edu/fastx_toolkit) were used to trim all reads to 150 bp length. Shorter sequences remaining after quality filtering and trimming, were removed with seqtk (http://github.com/lh3/seqtk). This was followed by interlacing of paired-ends with the FASTX-Toolkit. We then randomly selected 2× 1M, 2× 5M, and 2× 10M reads representing different genome coverages ranging from 0.02× to 2.04×. For read clustering, we used the Galaxy web interface of the RepeatExplorer pipeline2 [38, 39] with the parameters -l 39 (minimal overlap of clustering) and -o 30 (minimum overlap for assembly), considering the different genome coverages. The generated read clusters were further assigned to the retrotransposon lineages according to their similarity to reference elements in REXdb [28] and by RepeatMasker (http://www.repeatmasker.org, [40]). Based on the cluster graph shape, protein domains and repeat masker hits, each cluster was manually annotated and assigned to superclusters from the RepeatExplorer run of the highest genome coverage, which was expected to generate the highest number of full-length elements.

### Construction and classification of full-length LTR retrotransposons

To reconstruct full-length Ty1-*copia* and Ty3-*gypsy* retrotransposon sequences, several steps were followed: First, clusters and superclusters belonging to the individual retrotransposon types were identified based on their graph shape and protein domain hits. Second, contigs from the identified clusters were imported into Geneious Prime 2019, hereafter called Geneious (http://www.geneious.com, [41]). Third, we used the longest contig sequence as reference and mapped the remaining contigs against it using the Geneious mapper tool (with interation parameter = 10). We extracted the consensus sequence from the mapped contigs. Afterwards, short reads from the respective cluster were mapped against the consensus contig sequence to derive the final consensus sequence. If the identified LTR retrotransposon sequence was represented by only a single RepeatExplorer cluster, the consensus sequence was directly used for the reconstruction of the full-length element. However, if the LTR retrotransposon was split across multiple clusters, additional steps were considered for the reconstruction of the full-length element: These steps included iterative multiple sequence alignments of the consensus sequences of all clusters belonging to a single supercluster in Geneious. The parameters for the Geneious alignment tools were: automatically determine direction, alignment type = global alignment, cost matrix = 65 % similarity (5.0/-4.0), gap open penalty = 12, gap extension penalty = 3, refinement iterations = 2. Then, short read sequences from the respective supercluster were mapped against the reconstructed consensus sequence to derive the final full-length sequences using the Geneious mapper tool (with iteration parameter = 10) within Geneious.

The reconstructed full-length sequences were searched for the typical structural features of LTR retrotransposons, namely protein domain sequences, LTR regions, PBS, PPT, and internal repetitions. Different software and databases were used for these purposes: the LTR_Finder web server ([42]; http://tlife.fudan.edu.cn/tlife/ltr_finder/) was applied to predict common structural features along the reconstructed LTR retrotransposons, using the default parameters except “predict PBS by using tRNA database”, which was variable for different elements. Dotplot analyses were performed using the default parameter of the EMBOSS dottup tool integrated in Geneious [41, 43]. These dotplots showed the positions of LTRs as short diagonals in the upper right and lower left corners. Internal tandemly repeated structures lead to regular patterns of short lines in the dotplot. Furthermore, the REXdb database underlying RepeatExplorer’s protein domain finder tool (default parameters) was used to identify TE protein domains along the query elements and to predict their position and direction [38, 39]. Finally, the initial results from the protein domain finder tool were quality-filtered using the default parameters. For the Tandem Repeat Finder (TRF) tool the following parameters were used: alignment (match, mismatch, indel=2,7,7; 2,5,7; 2,5,5; 2,3,5), minimum alignment score to report repeats (20-60), and minimum period size (10-300) [44]. The TRF output was manually inspected, compared to the dotplot findings, and recorded for different features like period size, consensus sequence, number of repeat units, and position within the full-length LTR retrotransposon.

The online databases used to characterize and classify each full-length sequence were The Gypsy Database (GyDB) [25], Repbase [27], and REXdb database [28]. Repbase was used from the web-based software censor (https://www.girinst.org/censor/index.php) and GyDB from the online server (http://gydb.org/index.php/Main_Page). The REXdb database is integrated in RepeatExlorer2 (https://galaxy-elixir.cerit-sc.cz/).

The reconstructed full-length sequences were named according to the conventions and deposited in the Repbase database. Detailed information is summarized in Supplementary File 1 and Tables S1.

### Assignment of retrotransposon lineages and clades

A comparison against a database of well-known protein domains serves as the basis of a detailed retrotransposon classification. For this, protein sequences of reverse transcriptases (RTs) and, if applicable, chromodomains (CHD) were extracted from the publicly available genome assembly for *V. macrocarpon* (GenBank assembly accession: GCA_000775335.1) using the protein domain finder tool DANTE implemented in the RepeatExplorer Galaxy web-interface [38, 39]. DANTE was used with the quality filtering (minimum identity = 0.3; minimum similarity: = 0.4; minimum alignment length = 0.8; interruptions = 3). The program output 2149 sequences which were then clustered with a 90 % similarity threshold using CD-HIT ([45] http://weizhongli-lab.org/cd-hit). RT and CHD amino acid sequences were aligned with MAFFT [46] within Geneious (parameters: algorithm = auto [selects appropriate strategy from L-INS-I, FFT-NS-i and FFT-NS-2 according to data size], scoring matrix = “200PAM/K=2”, gap open penalty = 1.53, offsetvalue = 0.123). Pairwise distance matrix from the RT alignment was calculated in Geneious for boxplot visualizations, which were created with R graphics [47].

The resulting multiple sequence alignments of RT and CHD sequences were visualized as a dendrogram using the approximate maximum likelihood estimation algorithm of the FastTree tool [48] and the Randomized Axelerated Maximum Likelihood RAxML method [49] implemented in Geneious (default parameters). This allowed assigning the detected LTR retrotransposons to published lineages and clades according to their encoded RT and CHD sequences.

### Analysis of *cis*-regulatory element associated with the 5’ LTR of the full-length LTR retrotransposons

We extracted the 5’ LTRs of the identified Ty1-*copia* and Ty3-*gypsy* full-length elements. Ambiguous nucleotides from each of the 5’ LTR sequences were replaced by an N using the software “Sequence Manipulation Suite” [50] (http://www.bioinformatics.org/sms2/filter_dna.html). For the identification of regulatory sequence motifs, 5’ LTR sequences were individually searched against the Plant Care database [51] (http://bioinformatics.psb.ugent.be/webtools/plantcare/html), a database on *cis*-acting regulatory elements in plants.

### All-against-all comparison of full-length LTR retrotransposon sequences

To detect the sequence similarity within lineages, and also for fine-scale classification of the phylogenetically closely related sequences, all reconstructed *V. macrocarpon* full-length LTR retrotransposons were subjected to an all-against-all comparison together with publicly available reference sequences (Table S1). Representative reference sequences for each LTR retrotransposon lineage from different plant genomes were downloaded from GyDB 2.0 [25], (Table S2). The 5’ LTRs from these lineages were extracted using Geneious [41]. LTR regions were aligned in Geneious and a distance matrix of pairwise LTR identities was exported. For each lineage, an all-against-all sequence comparison was created using the Python-based software FlexiDot [52]. The FlexiDot parameters were: word size of 26 (-k 26) with 5 mismatches allowed (-S 5). In the dotplot, pairwise sequence identities of the 5’LTRs were printed and shaded.

### Transcriptome analysis

To assess retrotransposon transcription, publicly available *V. macrocarpon* paired-end mRNA-seq data (Illumina GAIIx; 63.6 million 100 bp reads) from leaves and shoot tips of cranberry cultivar ‘Ben Lear’ (accession number CNJ99-125-1) of BioProject PRJNA246586 were used [10]. All reads were quality-filtered using the FASTX-toolkit (http://hannonlab.cshl.edu/fastx_toolkit/) within Galaxy [53] with the parameters “quality cut-off” = 10, “percent above cutoff” = 95. Adapter trimming was performed using the BBDuk Geneious plugin [41, 54] with default parameters. About 36.1 million quality-filtered paired-end mRNA reads were used for read mapping against the reconstructed *V. macrocarpon* full-length elements of the Ty1-*copia* and the Ty3-*gypsy* superfamily. Transcriptome reads were mapped with the ‘Map to reference’ tool in Geneious [41] with the parameters: “mapper” = Geneious for RNA-Seq; “sensitivity” = medium-low sensitivity; “span annotated mRNA introns”. The graph coverage of the mapped reads was exported and the transcriptome proportion was calculated the counted reads in Excel. A scatter plot visualization was created to compare the genome proportion (derived from the RepeatExplorer analysis) and the transcriptome proportion of the representative full-length Ty1-*copia* and Ty3-*gypsy* LTR retrotransposons [47].

### Data availability

Full-length retrotransposon sequences and clustered core RT and CHD sequences from this analysis were deposited in the GDV database with the accession number GDV19002 (https://www.vaccinium.org/publication_datasets).

## RESULTS

### Composition of the *V. macrocarpon* retrotransposon landscape

The graph-based clustering with the RepeatExplorer2 pipeline reveals that about 91% of cranberry genome is repetitive (Fig. S1, [35]). According to the automated read cluster classification, only 49% of total repeat reads were assigned to known repeat types. Out of those, LTR retrotransposons constitute the major portion (44%), followed by DNA transposons (11%), non-LTR retrotransposons (6%), rDNAs (3%), satellite DNAs (0.06%) and other repeat fractions (Fig. S1, [35]). The *in silico* analysis shows that full-length LTR retrotransposons constitute a considerable portion of cranberry genome of 4.25 % Ty1-*copia* and 8.41 % Ty3-*gypsy* retrotransposons (Fig. S1; Table S1).

For a comprehensive overview of the LTR retrotransposon population, we identified the reverse transcriptase (RT) domains in the *V. macrocarpon* reference genome assembly. We queried the *V. macrocarpon* assembly and identified 181 Ty1-*copia* and 294 Ty3-*gypsy* RT instances. The highest number of RT sequences belonged to the Ale- (n = 110, Ty1-*copia*) and Tat- (n = 184, Ty3-*gypsy*) types (Table S3).

Maximum-likelihood analyses allowed us to verify the classification of the reference full-length elements for both superfamilies. For the Ty1-*copia* retrotransposons, we noticed a close relationship between five well-defined lineages: Ikeros, Bianca, Tork, TAR, and Angela. A close relationship of Ivana and SIRE sequences was also observed. Although Alesia only harbors two members, it forms a single, well-separated branch. Finally, the 110 Ale RT sequences are polyphyletic and strongly diversified (Fig. 1A). In the superfamily Ty3-*gypsy* (Fig. 2), the non-chromoviruses and chromoviruses are similarly organized. The non-chromovirus retrotransposons of the Tat type were the most abundant and the most diversified. The chromovirus lineage produced four branches representing the clades CRM, Reina, Galadriel, and Tekay (Fig. 2A), with a clade of non-chromovirus Athila as sister.

**Fig. 1.**
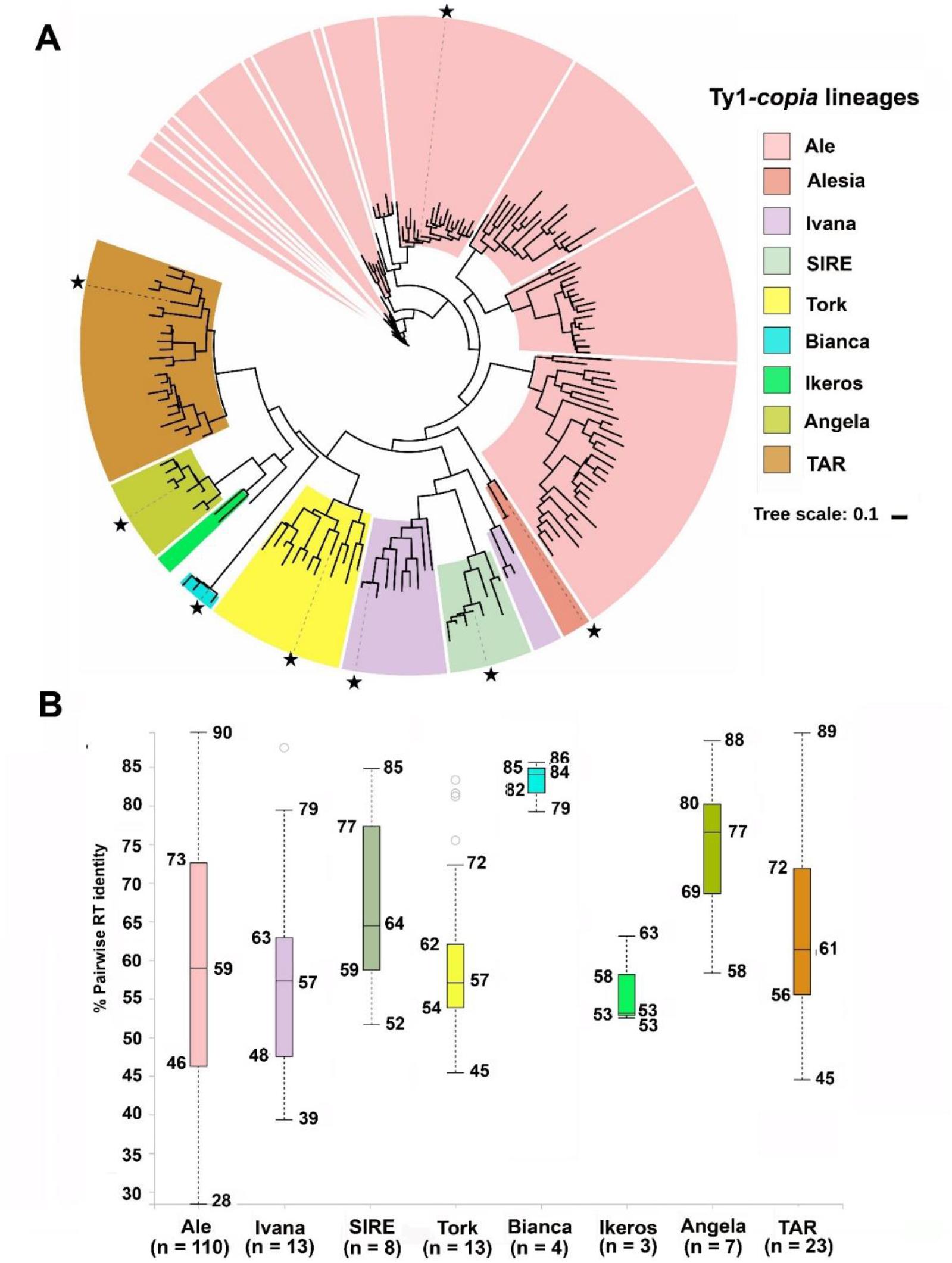
Diversity of the different Ty1-*copia* lineages in the *V. macrocarpon* genome. A) The dendrogram was derived from 181 Ty1-*copia* RT amino acid sequences calculated with the approximate Maximum-Likelihood method using the FastTree tool (Price *et al*. 2010) [45]. Lineages are color-coded (see internal legend). The stars indicate the position of the representative reconstructed full-length elements from *V. macrocarpon* genome shown in Fig. 3A. B) The boxplots illustrate the pairwise sequence identities of amino acid sequences of all Ty1-*copia* RTs. The lineage Alesia was excluded from the boxplots as only a single copy was detected in the *V. macrocarpon* genome. Colors in the boxplots correspond to panel (A). For each lineage, the total number of RT sequences (n) is provided, which includes the reference sequence.

**Fig. 2.**
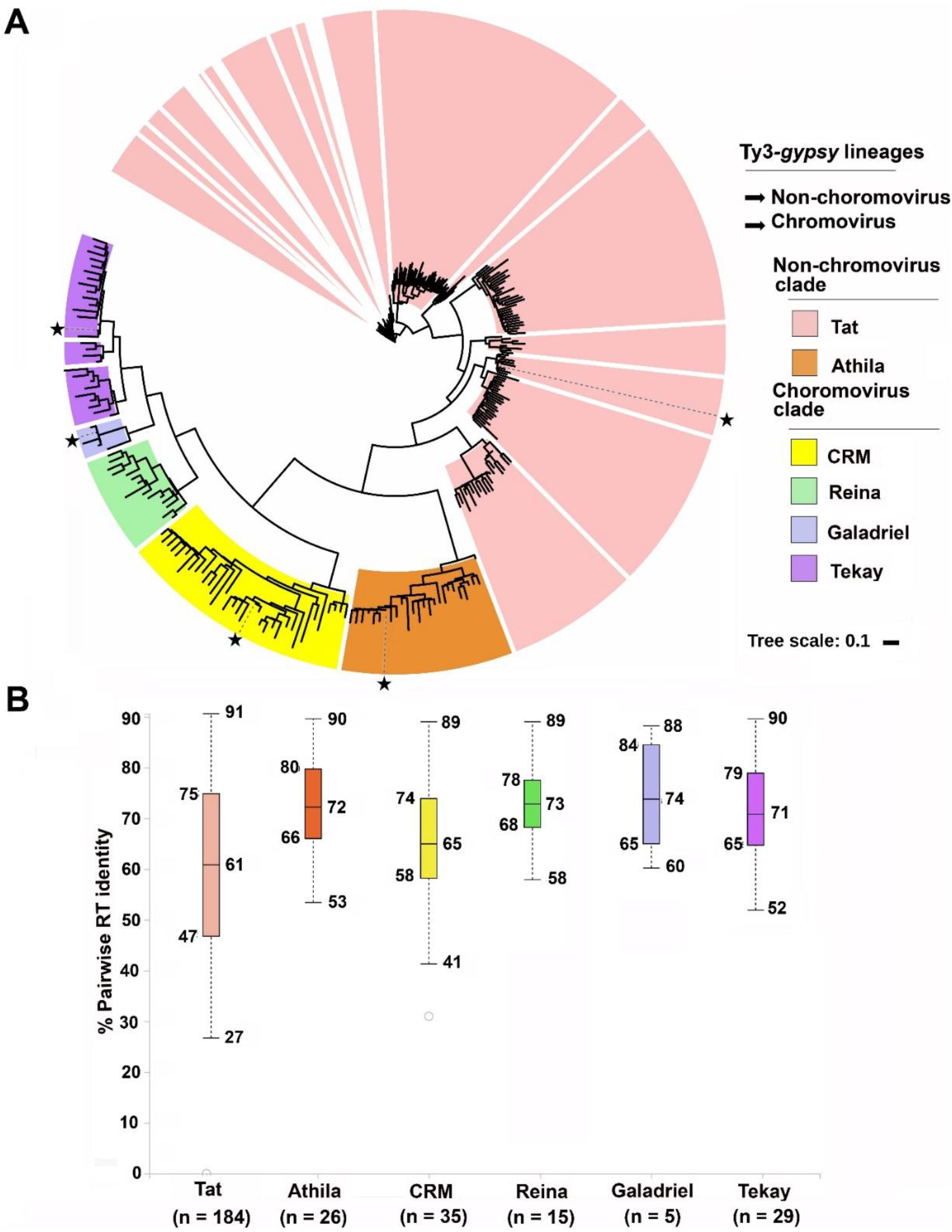
Diversity of the different Ty3-*gypsy* lineages in the *V. macrocarpon* genome. A) The dendrogram was derived from 294 Ty3-*gypsy* RT amino acid sequences calculated with the approximate Maximum-Likelihood method with the FastTree tool [45]. Clades are color-coded (see internal legend). The stars indicate the position of the representative for the reconstructed full-length elements from *V. macrocarpon* genome showed in Fig. 4A. B) The boxplots illustrate the pairwise sequence identities of amino acid sequences of all Ty3-*gypsy* RTs. Colors in the boxplots correspond to panel (A). For each clade, the total number of RT sequences (n) is provided, which includes the reference sequence

The separation of the chromoviruses into clades could be verified by reconstructing a dendrogram based on their signature chromodomain (CHD) sequences (Fig. S2). Although full-length Reina elements were not found in the RepeatExplorer-based reconstructed retrotransposon sequences (Table S4), the CHD search along the genome assembly of the same *V. macrocarpon* accession, reveals that CHD protein sequences are indeed present from all four chromovirus clades, i.e., CRM, Galadriel, Tekay, and Reina (Fig. S2).

Although branch lengths in the dendrogram provide information on the relative divergence between retrotransposon groups, the boxplot visualizations allow a direct comparison of the respective RT diversities (Figs. 1B, 2B). For this, pairwise identities from the RT amino acid multiple sequence alignments were utilized. In Ty1-*copia* lineages, Ale RT sequences have the highest diversity, as they exhibit the widest spread with a maximum of 90 %, a minimum of 28 %, and a median pairwise identity of 59 %. The lowest median identity (57 %) is observed for Ivana, whereas the highest conservation is observed for Bianca with a median pairwise identity of 84 % (Fig. 1B). Generally, Ty1-*copia* RT sequences are more diverse than those from Ty3-*gypsy*. Regarding the median RT identity, the most diverse clade in the Ty3-*gypsy* superfamily is Tat (61%), whereas Reina is the least diversified (73 %). The remaining clades of the Ty3-*gypsy* superfamily (Athila, CRM, Galadriel, and Tekay) have similarly diverse RT sequences, with median RT identities ranging between 65 % and 74 % (Fig. 2B).

### Structural diversity of Ty1-*copia* and Ty3-*gypsy* LTR retrotransposons

Based on the clustered short reads, we reconstructed and characterized 26 Ty1-*copia* and 28 Ty3-*gypsy* full-length retrotransposons, representative for 54 repeat families in the genome of *V. macrocarpon* (Table 1). The majority of these full-length elements (40 out of 54) was represented by a single, circular RepeatExplorer cluster in our analysis (Fig. S3). Reads from the remaining Ty1-*copia* and Ty3-*gypsy* elements were split into smaller clusters. Due to overlapping read pairs, we were able to connect these smaller clusters to superclusters spanning retrotransposons of full length (Table S1). Full-length consensuses were reconstructed from eight Ty1-*copia* lineages (Ale, Alesia, Bianca, Ivana/Oryco, TAR, Angela, Tork, and Sireviruses/Maximus, the last being incomplete) and three Ty3-*gypsy* lineages (non-chromoviruses Tat and Athila, and chromoviruses). Typical structural features, including sequence lengths, coding domains and conserved sequence motifs, were analyzed in detail to understand their structural diversity (Figs. 3, 4; Fig. S4; Table 1; Table S5).

**Table 1.**
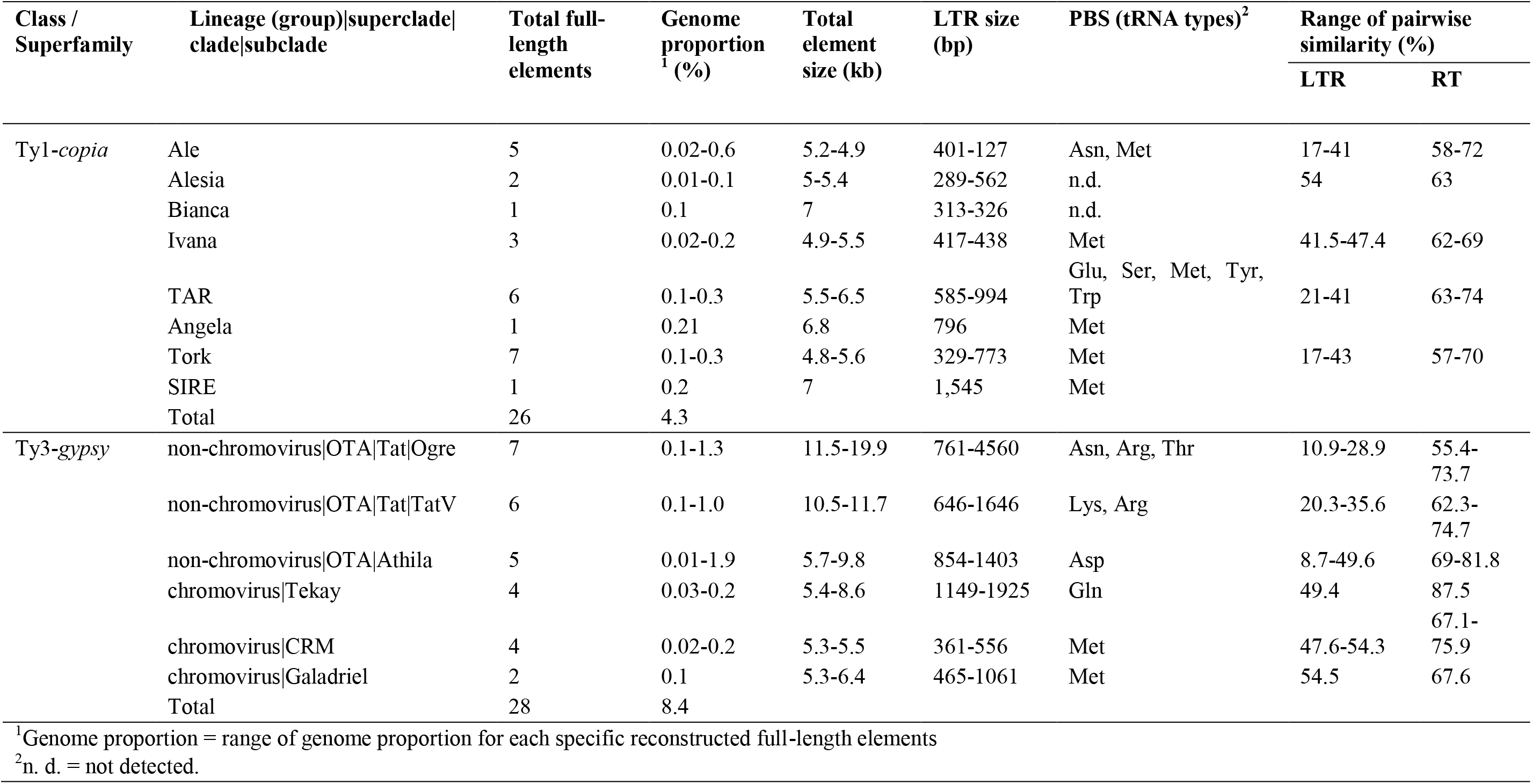
General structural features and repeat compositions of full-length LTR retrotransposons. Classification of the in silico retrotransposons was performed according to Neumann *et al.* 2019 [25].

**Fig. 3.**
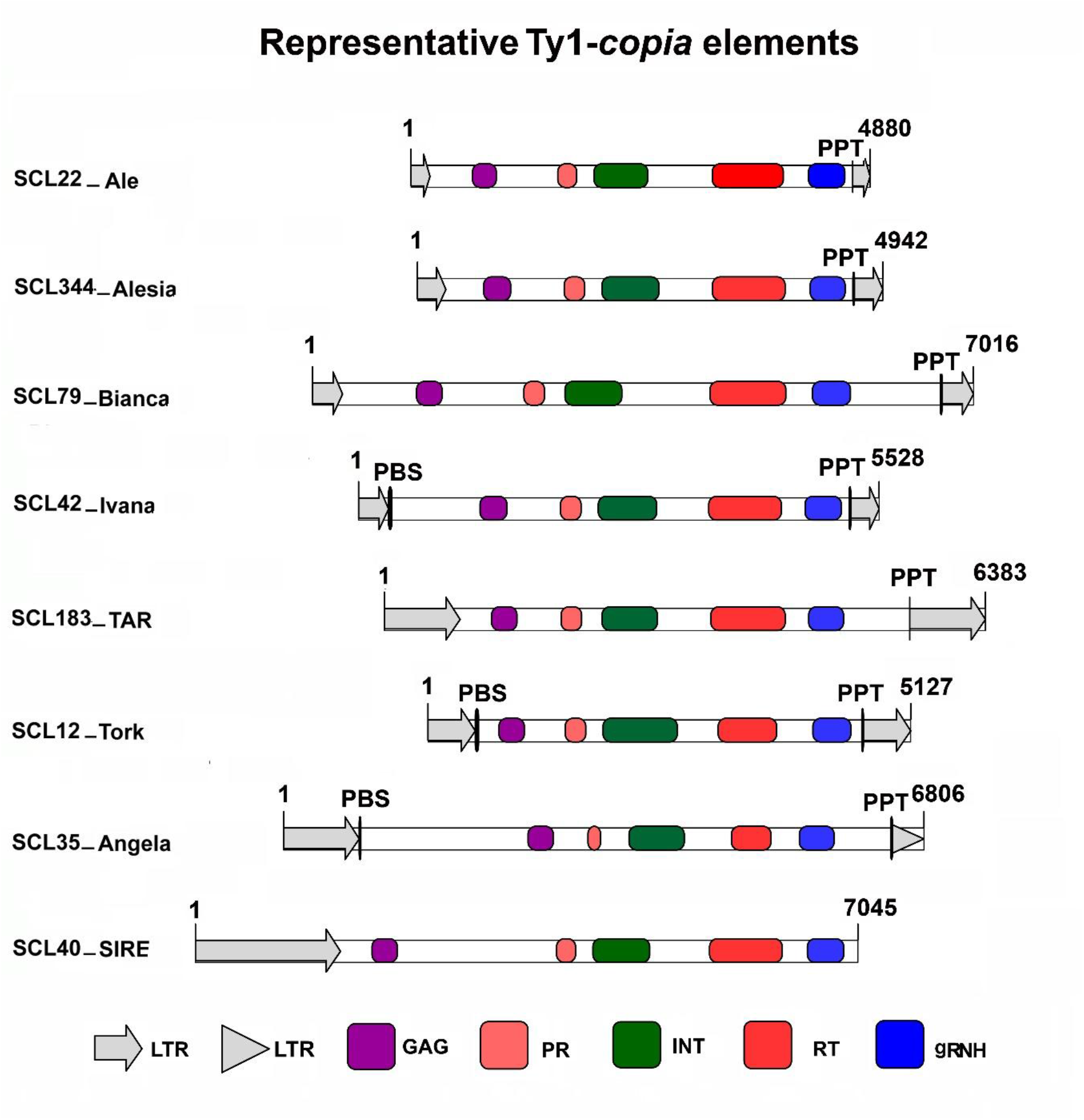
Structural features of representative *in silico* elements of Ty1-*copia* LTR retrotransposons in the *V. macrocarpon* genome. Long terminal repeats (LTRs) are represented as grey open arrows (intact LTR) and grey arrowheads (truncated LTR). Only the SCL40_SIRE consensus is incomplete and lacks the 3’ LTR. Black vertical lines adjacent to the LTRs represent primer binding site (PBS) and polypurine tracts (PPT), while thickness is according to length of this region (Table S2). Structural features of *gag* domain and the four genes of the *pol* domain are depicted as color-coded boxes: GAG = *gag* domain, PR = protease, RT = reverse transcriptase, gRNH = *gypsy*-type RNase H, INT = integrase.

**Fig. 4.**
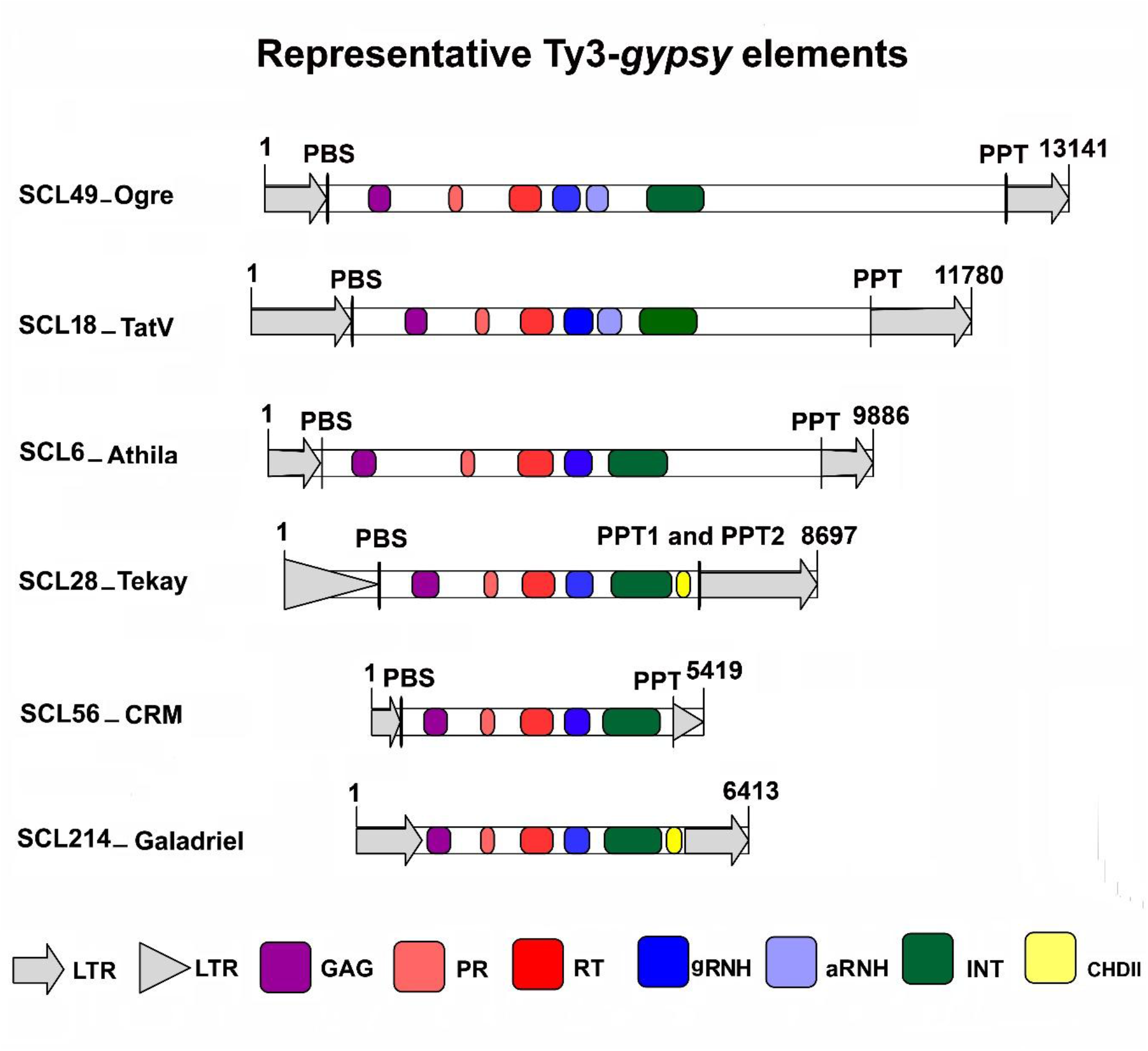
Structural features of full-length representative consensus elements of Ty3-*gypsy* LTR retrotransposons in the *V. macrocarpon* genome. Long terminal repeats (LTRs) are represented as grey open arrows (intact LTR) and grey arrowheads (truncated LTR). Black vertical lines adjacent to the LTRs represent primer binding site (PBS) and polypurine tracts (PPT), while thickness is according to length of this region (Table S2). Protein domains encoded in *gag* and *pol* are shown within the LTRs border. Structural features of the *gag* and *pol* genes are depicted as color-coded boxes: GAG = *gag* gene, PR = protease, RT = reverse transcriptase, gRNH = gypsy RNase H, aRNH = archaeal RNase H, INT = integrase, CHDII = chromodomain II.

To illustrate the structure of the individual Ty1-*copia* lineages, we selected eight representative elements (Fig. 3; Fig. S4). Among all the Ty1-*copia* lineages, we reconstructed seven full-length Tork, five Ale, two Alesia, one Bianca, three Ivana/Oryco, and one Sireviruses/Maximus (Table 1; Table S5): The longest consensus element belongs to the latter group and is about 7 kb, it is partially incomplete, containing only the 5’ LTR (1545 bp). Other lineages have retrotransposon lengths ranging between 4.8-6.8 kb with LTR lengths between 127-994 bp (Table 1; Table S5). While the highest genome proportion among the reconstructed Ty1-*copia* elements belongs to the elements SCL22_Ale (0.57 %), the lowest genome proportion is observed for SCL344_Alesia (0.01 %) (Table S5). Although all full-length *in silico* Ty1-*copia* elements contain uninterrupted *gag* and *pol* protein domains, some characteristic retrotransposon features were absent. For example, while most of the *in silico* elements that represent Ale, Angela and Tork retrotransposons harbored the PBS, PPT, and both 5’ and 3’ LTRs, one of these structural motifs was missing in the representative *in silico* sequences of Bianca, Ivana/Oryco, TAR, and Sireviruses/Maximus (Table 1; Table S5). The most common PBS-type detected in Ty1-*copia* lineages corresponds to the Met-type. For Ale and TAR retrotransposons, however, a PBS of the Asn/Met-type (Ale) or Glu/Ser/Met/Tyr/Trp-type (TAR) was more common (Table 1; Table S5).

Six reference elements were selected to represent the structural variability of each Ty3-*gypsy* lineage (Fig. 4). The most diversified and abundant Ty3-*gypsy* elements in the *V. macrocarpon* genome belong to the Tat group (Table 1). A total of 13 full-length elements have been characterized, with genome proportions ranging between 0.12 and 1.34 % (Table 1). In contrast, the other two groups only contribute 0.01-1.92 % (Athila) and 0.02-0.23 % (chromoviruses) to the genome (Table 1; Table S5). Although *gag* and *pol* genes were detected for all full-length elements, structurally complete elements with all motifs typical for LTR retrotransposon (PBS, PPT and both 3’ and 5’ LTR regions) were mostly present in the Galadriel clades (chromoviruses). Most elements of the other groups (chromovirus Tekay, CRM and non-chromoviruses Tat and Athila) lack at least one of these structural motifs. Moreover, some Ty3-*gypsy* elements have additional protein domains as compared to Ty1-*copia* elements. For instance, the *pol* genes of the non-chromovirus Tat clade have dual ribonuclease H domains (gRNH and aRNH), whereas the chromoviruses harbor an additional chromodomain (CHD) region (Fig. 4; Table S5). A canonical CHD was only found for the elements of the Tekay and the Galadriel clades, but surprisingly not within the elements of the chromovirus CRM clade (Fig. 4; Fig. S4; Table S5). SCL16_Ogre is the longest Ty3-*gypsy* Tat element, with a total length of 19 kb and 1251-1326 bp long LTRs. In contrast, SCL80_CRM is the shortest element (5.2 kb) with LTRs ranging between 361-556 bp (Table S5). A Met-type PBS was only found in the CRM and Galadriel chromoviruses, whereas other Ty3-*gypsy* elements were more variable regarding their PBS (Table 1).

### Cranberry retrotransposons are often associated with short tandem repeats

Although tandem repeats, i. e. satellite or ribosomal DNAs, represent major repetitive genome fractions that are distinct from the more dispersed transposable elements, tandemly repeated motifs can also be associated with retrotransposons. In *V. macrocarpon*, we detected tandem repeats in the reconstructed full-length Ty1-*copia* and Ty3-*gypsy* retrotransposon sequences (Table 2; Table S6). These tandem repeats were either localized within the 3’ or the 5’ LTRs, within the untranslated regions (UTRs) adjacent to either LTR, or in some cases within the UTRs adjacent to the *gag* gene. Tandem repeats found in different retrotransposons vary in sequence composition, period size, and number of repetitions (Table 2; Table S6). For the Ty1-*copia* superfamily, we identified tandem repeats in all lineages, embedded in either the LTRs or UTRs, with consensus period lengths ranging from 2 to 49 bp with about 2 to 8 repetitions (Table 2; Table S6).

**Table 2:** Summary of tandem repeats and *cis*-regulatory elements associated with the reconstructed full-length LTR-retrotransposon sequences.

| **Lineage**<br>**/clade**<br>**/subclade** | **Location of tandem repeats** | **Consensus**<br>**period size** | **Number of**<br>**repetitions** | **Total number of different *cis*-regulatory**<br>**motifs identified in 5' LTR region** |
| --- | --- | --- | --- | --- |
| Ale | 5' and 3' LTR, 5' UTR | 2-48 | 1.9-7.9 | 10 |
| Alesia | 5 LTR, 5' UTR | 30 | 2.2-4.9 | 5 |
| Bianca | 5'UTR | 25 | 2.8 | 2 |
| Ivana | 5'LTR | 27 | 2.9 | 8 |
| TAR | 5'LTR | 10-17 | 1.9-4.4 | 19 |
| Angela | 5'LTR | 18 | 5.4 | 9 |
| Tork | 5'and 3' LTR, 5' UTR | 11-49 | 2-6.8 | 19 |
| SIRE | 5' LTR | 15 | 2 | 12 |
| Ogre | 5' and 3' LTR, UTR adjacent<br>to the *gag* gene | 6-91 | 2-14.2 | 38 |
| TatV | 5' and 3' LTR, UTR | 15-93 | 2.1-15.1 | 24 |
| Athila | 3' LTR, UTR | 21-42 | 2-2.5 | 26 |
| Tekay | 5' and 3' LTR, UTR | 2-29 | 1.9-18.5 | 10 |
| CRM | 5' and 3' LTR, UTR | 17-48 | 1.9-4.5 | 8 |
| Galadriel | n.d. | n.d. | n.d. | 13 |
| n.d. = no tandem repeat detected |  |  |  |  |

In Ty3-*gypsy* LTR retrotransposons, tandem repeats were detected both in non-chromoviruses and chromoviruses, yet to a different extent. Non-chromovirus Tat and chromoviruses harbored tandem repeats both within the LTRs and UTRs, whereas non-chromovirus Athila had tandem repeats only embedded within the UTR adjacent to the *gag* gene (Table 2). Among all Ty3-*gypsy* LTR retrotransposons, Tat (including Ogre) harbor tandem repeats with the largest monomer size (up to 93 bp) and the highest number of repetitions (up to 15 times) (Table 2; Table S6).

### *Cis*-regulatory elements are often associated with LTR sequences

As LTRs frequently harbor *cis*-regulatory elements, our targeted search revealed that almost all reconstructed retrotransposons include typical promoter motifs, including TATA and CAAT boxes within their LTRs (Fig. S5A-B; Table 2; Table S7). However, as the LTR composition varies, number and position of these promoter sequences are variable. For instance, the LTRs of the Ty1-*copia* lineages Ale, Angela, TAR, Tork, Sireviruses/Maximus, and all Ty3-*gypsy* lineages contained multiple promoter motifs in different positions. However, the Bianca LTR only contains a CAAT box, lacking a TATA box. For Ivana, two out of three *in silico* elements (SCL42_Ivana and SCL310_Ivana) contain both promoter motifs in their LTRs (Fig. S5A-B; Table S7).

In addition to the promoter motifs, we detected other *cis*-regulatory elements for different LTR retrotransposons (Table S7). These regions are mostly related to light, hormone, and cell cycle responsiveness, as well as abiotic stress tolerance (Fig. S5A-B; Table S7). Among the eight representative sequences of the Ty1-*copia* lineages, most regulatory motif types were predicted for TAR (19), Tork (19), and Ale (10) (Table 2), whereas among the Ty3-*gypsy* elements, the Tat retrotransposons (including Ogre) harbor most variability with 38 distinct motifs.

### Sequence homogeneity between closely related elements

In an attempt to fine-scale the classification of the full-length retrotransposon elements of *V. macrocarpon*, closely related elements were subjected to an all-against-all sequence comparison with publicly available references to reveal sequence similarities among them (Table S2).

Overall, the *V. macrocarpon* full-length elements have higher similarity to other elements within their protein coding regions, contrasting the variability within their UTRs and LTRs (Fig. S6A-E). However, the levels of similarity for coding and non-coding regions vary depending on the lineage (Fig. S4). For instance, the RT similarities within the Ty1-*copia* and within the Ty3-*gypsy* groups ranged between 57-74 % and 55-88 %, respectively. In contrast, LTR similarities ranged between 17-47 % for Ty1-*copia* groups and between 11-55 % for Ty3-*gypsy* groups, respectively (Fig. 6A-E; Fig. S4; Table 1). Ale and Tat sequences are the most diversified, both within their RTs and their LTRs (Table 1).

### Transcriptome analysis reveals transcriptional activity of LTR retrotransposons

We investigated whether the identified full-length LTR retrotransposons are transcribed into RNA by analyzing the reference transcriptome database of *V. macrocarpon* from leaves and shoot tips (Fig. 5; Fig. S7; Table S8). We have mapped a total of 36.1 million transcript sequences against the 54 reconstructed Ty3-*gypsy* and Ty1-*copia* sequences of *V. macrocarpon.* Subsequently, we compared the genome proportion derived from the RepeatExplorer analysis against the transcriptome proportion for each individual element (Fig. 5).

**Fig. 5.**
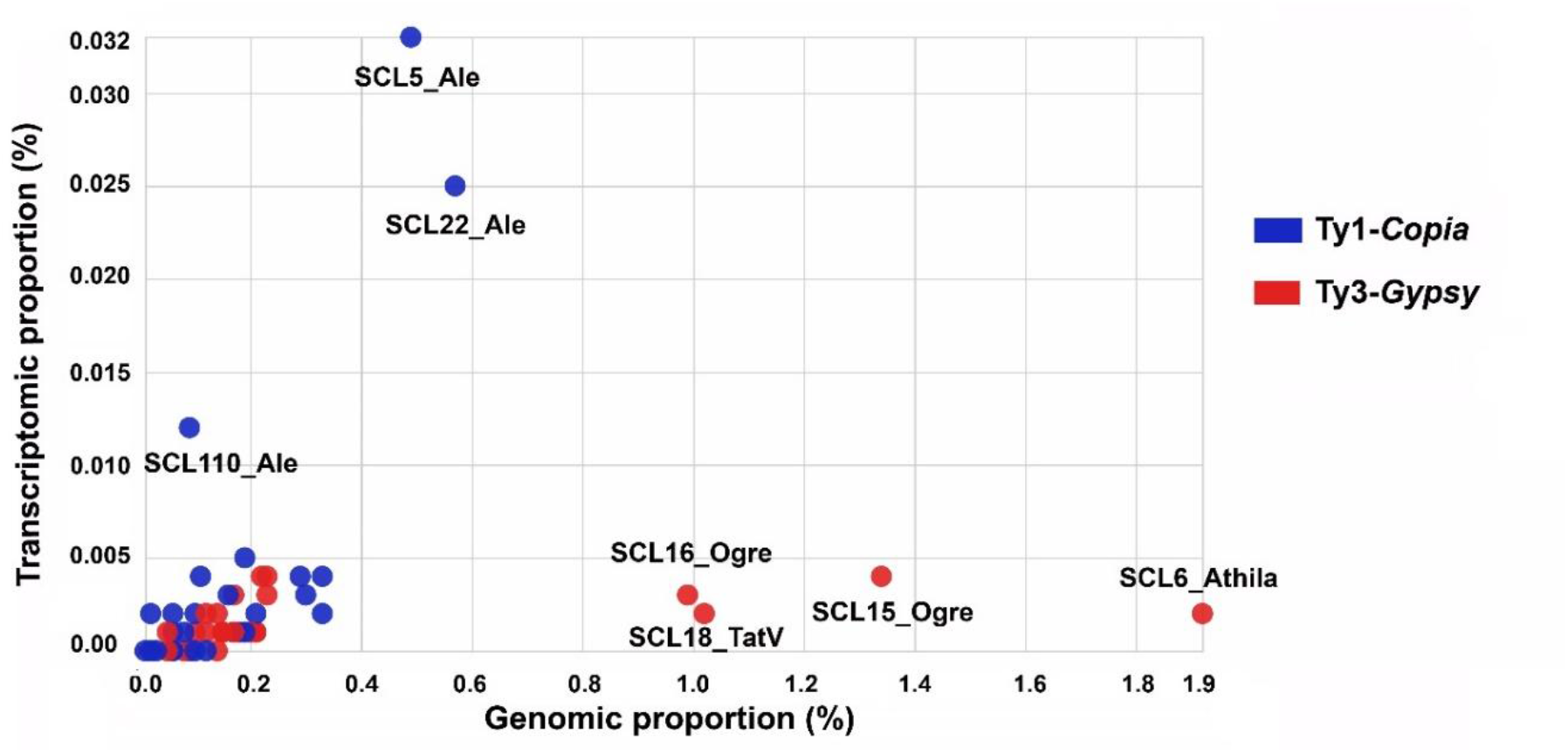
Comparison of genome and transcriptome proportions for the 26 Ty1-*copia* and 28 Ty3-*gypsy* full-length consensus retrotransposons from *V. macrocarpon*. The genome proportions of the reconstructed full-length Ty1-*copia* and Ty3-*gypsy* elements are plotted against their transcriptome proportions. Genome proportions were calculated from the RepeatExplorer output (Table S2). Transcriptome proportions were calculated from read mapping of the publicly available Illumina cDNA sequence reads of *V. macrocarpon* (accession number PRJNA246586). Major groups of the 54 elements are color-coded (see internal legend). Only the names of elements of the highest genome and/or transcriptome proportions are annotated. For details, see Table S2.

All the identified full-length elements are transcribed to a different extent, thereby contributing a total of 0.1 % by Ty1-*copia* retrotransposons and 0.03 % by Ty3-*gypsy* retrotransposons to the used *V. macrocarpon* transcriptome reads (Fig. 5; Fig. S7; Table S8).

Among all LTR retrotransposon lineages, Ale has the highest proportion of transcript sequences (Fig. 5; Table S8). In fact, the two reference elements SCL5_Ale and SCL22_Ale have both the highest genomic proportions of all 26 Ty1-*copia* representatives and the highest percentage of mapped RNA reads of all 54 reference LTR retrotransposons in *V. macrocarpon* (Fig. 5; Table S8). Nevertheless, the results also show that the percentage of mapped transcripts is dependent on the individual full-length elements and cannot be generalized across and within lineages. Nevertheless, the number of mapped transcript sequences is not proportional to their genomic abundance: most lineages, including Angela, Bianca, Ivana, TAR, Tork, and SIRE, have less representative transcript sequences as compared to their proportional genomic abundance (Fig. 5; Table S8).

Despite their higher genomic abundance, the transcription level of Ty3-*gypsy* retrotransposons is much lower (0.037%) than the transcription of Ty1-*copia* retrotransposons (0.105%): SCL15_Ogre, SCL27_Ogre, SCL28_Tekay, and SCL31_CRM generate most transcripts with similar counts (0.003-0.004 % of the total transcript sequences; Fig. 5; Table S8). Noteworthily, SC31_CRM has only a low genomic abundance (0.22%), but the highest number of transcripts (0.004%) as compared to other Ty3-*gypsy* elements.

We have also investigated the distribution of mapped transcript reads along the most transcribed full-length Ty1-*copia* and Ty3-*gypsy* elements to gain insight into the coverage of the different structural features. In Fig. S7, the most transcribed elements from each group and the distribution of mapped RNA reads are shown. The distribution of mapped transcript sequences along the different structural components of the LTR retrotransposons shows either a homogenous profile along the full-length element or an enrichment in the LTRs and UTRs. SCL5_Ale and SCL49_Tork have homogeneously mapped transcript sequences along their *gag*-*pol* genes and UTRs (Fig. S7). In contrast, SCL42_Ivana, SCL40_SIRE, SCL49_Ogre, and SCL31_CRM have the highest number of mapped transcript reads in the two LTRs. Moreover, SCL79_Bianca and SCL6_Athila have most transcripts mapped to the internal UTR located between the *gag* and *pol* genes as well as to the 3’ LTR (Fig. S7).

## DISCUSSION

Although a draft genome assembly for *V. macrocarpon* is already available [10], we are only starting to decipher its repetitive fraction. As we showed in our previous work that focused on satellite DNAs in *Vaccinium* genomes [36], repetitive DNAs are still the last genomic portions to be assembled completely. Nevertheless, as knowledge of repetitive DNAs is needed to understand *Vaccinium* genomes in their entirety, we extend our previous work to characterize the contribution of long terminal repeat (LTR) retrotransposons to the cranberry genome. As we base our analysis on a random sampling of Illumina reads, we avoid biases introduced by a potentially erroneous genome assembly [39]. Nevertheless, as this approach may neglect less abundant and more diverged repeats [55], all quantifications are likely underestimations.

We showed that nearly a quarter of the *V. macrocarpon* genome is made up of highly abundant, repetitive TE families, with over 91% of genome considered as repetitive [35]. This nearly doubles the estimate retrieved for the cranberry genome assembly (40%) [10, 11], and likely is a result of a systematic exclusion of repetitive regions from the genome assembly. Within the repetitive genome fraction, LTR retrotransposons are the most abundant TEs (44 % of the repetitive fraction), followed by DNA transposons (11 %), and non-LTR retrotransposons (6 %). The observed high abundance of LTR retrotransposons is typical for plants and compares well to other TEs in other species such as flax [56], apple [57] maize [58], and oats [59].

### Cranberry LTR retrotransposons are diverse in sequence and structure

Overall, in the *V. macrocarpon* genome, Ty3-*gypsy* elements are on average longer and contribute a higher genome proportion than Ty1-*copia* elements. This was also reported in other plants, such as *Avena sativa* [59], *Triticum* sp. [60], *Festuca pratensis* [61], and *Malus domestica* [57]. Nevertheless, opposite proportions are also possible, as in *Musa acuminata* and *Pyrus bretschneideri*, in which Ty1-*copia* retrotransposons outweighed Ty3-*gypsy* elements [62]. Variation in sequence lengths between these two groups is thought to reflect length differences in the non-coding rather than in the coding element regions, whereas the genome proportion depends on their copy number rather than on the mere retrotransposon length [63–65]. However, elements from Errantivirus/Athila and Ogre/Tat lineages from Ty3-*gypsy* were found to have both the highest genome proportions and the largest sizes in the *V. macrocarpon* genome. Besides the generally large lengths of Ogre/Tat retrotransposons in *V. macrocarpon*, the high number of RT sequences detected in this study also indicates a high copy number – both contributing to their high genomic abundance. Similarly, for the Ale lineage, we found the most RT instances and a higher genome fraction as compared to the other Ty1-*copia* lineages.

Characterization of *V. macrocarpon*’s RT sequences revealed that Tat and Ale are the most diversified groups among the Ty3-*gypsy* and Ty1-*copia* retrotransposons, respectively. By studying the pairwise RT identities, we observed that individual lineages diversify differently. This is in accordance with our previous report on the related diploid species *Vaccinium corymbosum* [35]. Species- and genus-specific lineage diversification and heterogeneity is reported by many authors for several species. For example, within Ty1-*copia*, Sirevirus/Maximus retrotransposons were highly diversified in sugarcane [64], in the Fabeae [65], and in maize [66], whereas Ale retrotransposons were found in highest diversity in *Chenopodium quinoa* [67] and *Populus trichocarpa* [68]. Within Ty3-*gypsy*, the Tat lineage was found to proliferate in sugarcane [64] and in the Fabeae [65], whereas the Athila clade in *Populus trichocarpa* [68], and the Tekay chromoviruses was for example more abundant in *Beta vulgaris* [69]. The overall abundance and diversification of the various LTR retrotransposon types also reflects the evolutionary relationships of the host species and may be similar within a group of closely related species [70, 71]. The environmental adaptation of a particular species may also be affected by the insertion and molecular diversification of these retrotransposons [16, 72].

Structurally, regarding the encoded protein domains, the reverse transcriptase (RT) is the most ancient domain and key to the dispersion of retrotransposons [17, 73]. The RT is present in all autonomous LTR retrotransposons, whereas the modular acquisition of other polyprotein domains is assumed to result from independent lineage-specific evolutionary histories [17, 74]. Therefore, structural differences among the lineages are considerable. This is particularly evident in the Tat and chromovirus lineages that harbor additional protein-coding domains. All members of the Tat lineage in *V. macrocarpon* harbor two ribonuclease H domains. Ustyantsev *et al.* (2015) [75] speculated that this duality of ribonucleases H might benefit the successful strand transfer, a property common to retroviruses that suffer natural selection pressure during proliferation.

Chromoviral integrases are typically marked by a chromodomain at their C-terminus, presumably with the ability to bind specific histone variants [69 76-79]. Therefore, chromoviruses may directly impact chromatin structure and function [79]. For the chromovirus clade Galadriel and Tekay, a chromodomain was detected both in the *V. macrocarpon* genome assembly and in the representative full-length elements from the read cluster analysis. In contrast, for the CRM clade, chromodomain regions were extracted from the genome assembly only, although both genome assembly and read cluster analysis were based on the same *V. macrocarpon* accession. Compared to the CHDs of the Tekay, Galadriel, and Reina clades, their identification within centromeric retrotransposons is more difficult, as they typically do not show structural similarities to type I and II cellular chromodomains and are highly variable even between families within a species [69, 78]. Nevertheless, this domain, also referred to as the CR motif or CHDCR, correlates with various sequence and structural features [28]. For example, the RTs of CRM elements tend to form a separate branch in the analysis of sequence relationship. Another feature is the position of the CHDCR, which is defined by its extension into the 3’ LTR. Although some of the reference elements of the CRM clade have incomplete LTR sequences, an extended *gag-pol* ORF into the 3’ LTR is recognizable. CR chromodomains of CRM elements were found to be mostly associated with the centromeric heterochromatin. These domains are likely responsible for the diversification, integrity, and stability in the functional centromeric heterochromatin through an RNA-mediated mechanism in different beet and grass species [69, 79–81].

The PBS is an important structural feature for LTR retrotransposon transcription located downstream of the 5’ LTR, with PBS type and length characteristic for a given plant LTR retrotransposon group [28]. The PBS is present in most of the *in silico* retrotransposons constructed for *V. macrocarpon*. Nevertheless, some consensuses lack a designated PBS, likely either an assembly or identification error, or the nature of the PBS itself in the studied species. In chromoviruses as well as in Ty1-*copia* retrotransposons, a PBS corresponding to the initiator methionine tRNA (Met-tRNA) was most common, similar to other plant genomes [28, 82]. In contrast, the PBS regions in other Ty3-*gypsy* retrotransposons were more diverse. The specific PBS site is not only significant for the transcription of LTR retrotransposons, but also for their post-transcriptional regulation through tRNA-derived small RNAs [82, 83]. Therefore, the acquisition of distinct PBS sequences in different retrotransposon lineages may point to individual evolutionary benefits.

The LTRs of the identified full-length Ty1-*copia* and Ty3-*gypsy* retrotransposons were enriched for different *cis*-regulatory elements, including putative promoters and sequences related to hormone responsiveness, cell development, and stress response. This indicates the relevance of these motifs for retrotransposon transcription and activity. In addition, the *V. macrocarpon* full-length elements harbored tandem repeats not only in their LTR region but also in their UTRs. Yet, none were found within coding sequences. The presence of *cis*-regulatory regions and tandem repeats in the LTR is common for LTR retrotransposons and can have significant impacts on their reverse transcription and proliferation [25, 55]. Moreover, *cis*-regulatory sequences could act as enhancers or repressors and thereby affect the expression of downstream genes as part of regulatory networks and pathways [55, 84]. Although still a matter of investigation, the origin of tandem repeats from different parts of the retrotransposons (i.e. LTRs or UTRs) with specific functions are reported for many organisms. An example are the centromeric tandem repeats that may have originated from different retrotransposons in maize [85], wheat [86], and potato [87], suggesting their importance for the formation of functional centromeres in these species. Moreover, the existence of short tandem repeats varying in length and sequence in UTR sequences near the PPT was also reported for many other plant genomes, such as legumes [63] and *Silene latifolia* [88]. Although the specific function of these short repeats is still debated, it is hypothesized that they could serve as a junction, connecting the *gag-pol* gene to the 3’ end. Thus, in evolutionary time-spans, they may serve as a seed for the generation of longer satellite DNA arrays, with the potential to acquire a structural function for the organization of chromosomes [88].

We found that the overall similarity among the full-length elements within each lineage varied. For example, Tat retrotransposons are diversified and have little overall sequence similarity within the UTR regions as was also reported for other organisms [63, 65]. The highly diverse UTRs and LTRs on clade level (see Fig. S6) may indicate their ancient origin and diversification in the *Vaccinium* ancestors [28]. LTRs and UTRs appear to evolve faster than coding regions due to different mechanisms, like unequal recombination or illegitimate recombination [63, 89–92].

### Cranberry LTR retrotransposons of the Ale-type are strongly transcribed in *V. macrocarpon* leaves and shoot tips

As expected, all *V. macrocarpon* LTR retrotransposons investigated showed at least a basal level of transcription, together making up about 0.13 % of the total transcriptome. According to [10], transcription of TEs in *V. macrocarpon* is quite low (about 4.3%) and in contrast to the highly transcribed protein-coding genes (about 83.59%). This is in line with our study, in which only the most abundant LTR retrotransposons were queried. A low transcription of retrotransposons compared to conventional protein-coding genes is a common finding in most plant species, including grasses [93], flax [56], and sugarcane [64]. The low transcription of full-length LTR retrotransposons indicates out that these sequences are rather strictly regulated in cranberry [10].

Transcriptional activity of full-length LTR retrotransposons is not correlated with their genome proportion in *V. macrocarpon*. Higher levels of expression than those found here for less abundant elements (such as Ale elements) have been reported in sugarcane [64], maize [58], poplar [94], *Arabidopsis arenosa* [95], and sunflower [96]. Overall, Ty1-*copia* sequences appeared to be more transcribed than Ty3-*gypsy* elements in *V. macrocarpon* genome being the highest transcription profiles (∼0.08%) those of the Ale full-length elements. In contrast, Athila elements have a higher genomic percentage (2.44%), but have fewer transcript reads assigned (0.004 %), even not covering the full reference element length. In fact, it is considered that Ale and Sireviruses are some of the most transcriptionally active Ty1-*copia* lineages in plants [58, 64, 66, 95–97]. In cranberry, for Ale retrotransposons, this also seems to hold true, whereas for Sireviruses, fewer genomic and transcriptomic proportions were detected. Nevertheless, both Ale and Sireviruses are preferentially accumulated in the euchromatic region of the genome in different species [58, 95, 96] and hence could affect gene regulatory pathways.

In contrast, retrotransposon heterochromatization and fragmentation may decrease element transcription. We know from other plant genomes that some lineages are more prone to truncation and heterochromatic burial, e.g. Athila/Errantiviruses or even the related pararetroviruses in sugar beet [98, 99]. Similar tendencies may impact the TE landscape in *macrocarpon*, yielding lineages that are less transcribed than others, e.g Athila/Errantiviruses and Sireviruses. Transcriptional activity of TEs is generally dependent on several factors and may be correlated with development [100], (epi)genetic regulation [15, 16, 64, 101], and environmental adaptation [15, 95, 100]. Therefore, TE transcription can depend on the genomic neighborhood, the presence of *cis*-regulatory motifs, tissue types, developmental states, and environmental effects [100, 101]. Here we observed considerable differences regarding the genome and transcriptome abundances of repetitive sequences in *V. macrocarpon*, which imply that the mechanisms of transcriptional regulation vary depending on the LTR retrotransposon.

Regarding their applicability, the retrotransposons identified here may provide suitable targets for the development of molecular markers in assisted breeding programs [19–23]. We suggest targeting retrotransposons that are most abundant and for which high transcription implies closeness to genes. Hence, we would suggest Ale retrotransposons within the Ty1-*copia* superfamily, as well as Athila retrotransposons within the Ty3-*gypsy*. Both are abundant in genomic and transcriptomic databases, and thus may provide many primer binding sites and likely are situated in the more openly packed euchromatin.

### Conclusion

Using short reads and the sequence assembly, we have provided an exhaustive overview of the LTR retrotransposon landscape in the cranberry genome. We have detected all major LTR retrotransposon lineages, with some showing association to short tandemly repeated motifs. Considering their high genomic abundance and transcriptional activity, we suggest that Ale and Athila TEs likely represent the most useful targets to survey TE-derived polymorphisms across genotypes.

## Conflict of interest

The authors have no conflict of interest to report.

## Acknowledgments

This work was supported by the Scientific and Technological Research Council of Turkey (TUBITAK) TUBITAK-2215 PhD scholarship and the Scientific Research Projects Unit (BAP) of Niğde Ömer Halisdemir University (FEB 2017/18 DOKTEP). This work also acknowledges the financial support by the Jagannath University, Bangladesh, in the form of study grants as well as the Georg Forster fellowship of the Alexander von Humboldt Foundation awarded to NS.

## Abbreviations

LTR - long terminal repeat; TEs - transposable elements; PR - protease; gRNH - RNase H; RT- reverse transcriptase; INT - integrase, PBS - primer binding site; PPT - polypurine tract; CHD – chromodomain; UTR – untranslated region; DNA - deoxyribonucleic acid; RNA - ribonucleic acid.

## Notes

https://www.vaccinium.org/vaccinium_downloads/vaccinium_projects/GDV19002-V_macrocarpon_LTRs/

### Competing Interest Statement

The authors have declared no competing interest.

