## Supplementary Figures 1-7 for "Genome-wide analysis of long terminal repeat retrotransposons from the cranberry *Vaccinium macrocarpon*"

**This PDF file includes:**

Figures S1 to S7

**A RepeatExplorer output without any treatments from *Vaccinium macrocarpon* genome**

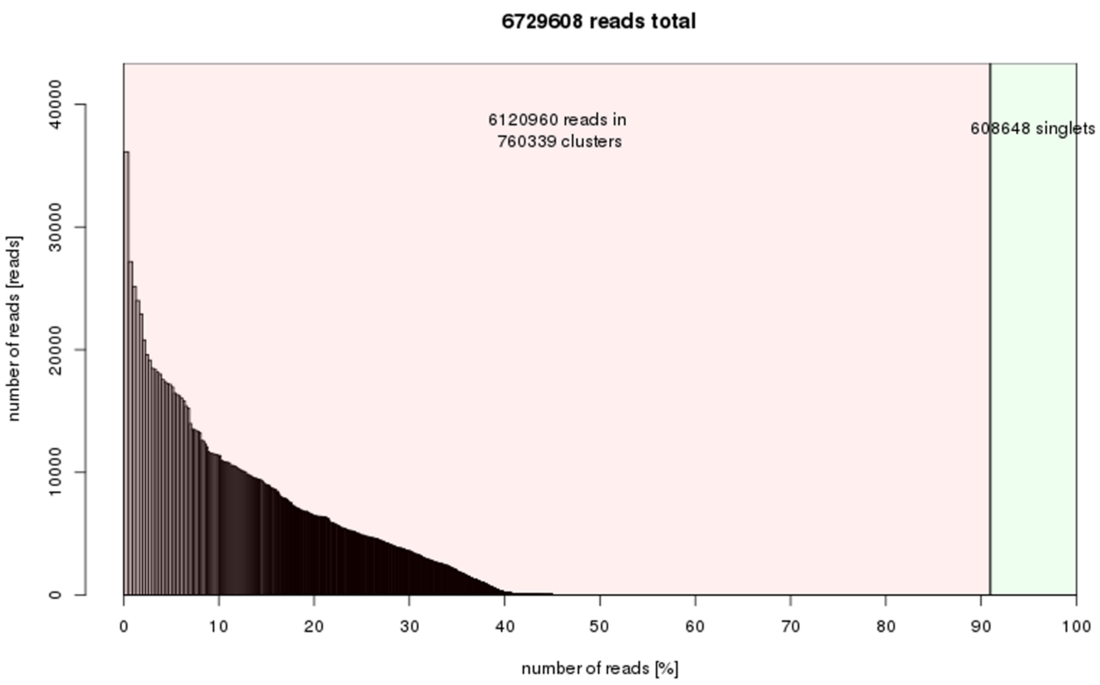


**B Assignment of repeat types to the most abundant repeat clusters in *Vaccinium Macrocarpon* genome**

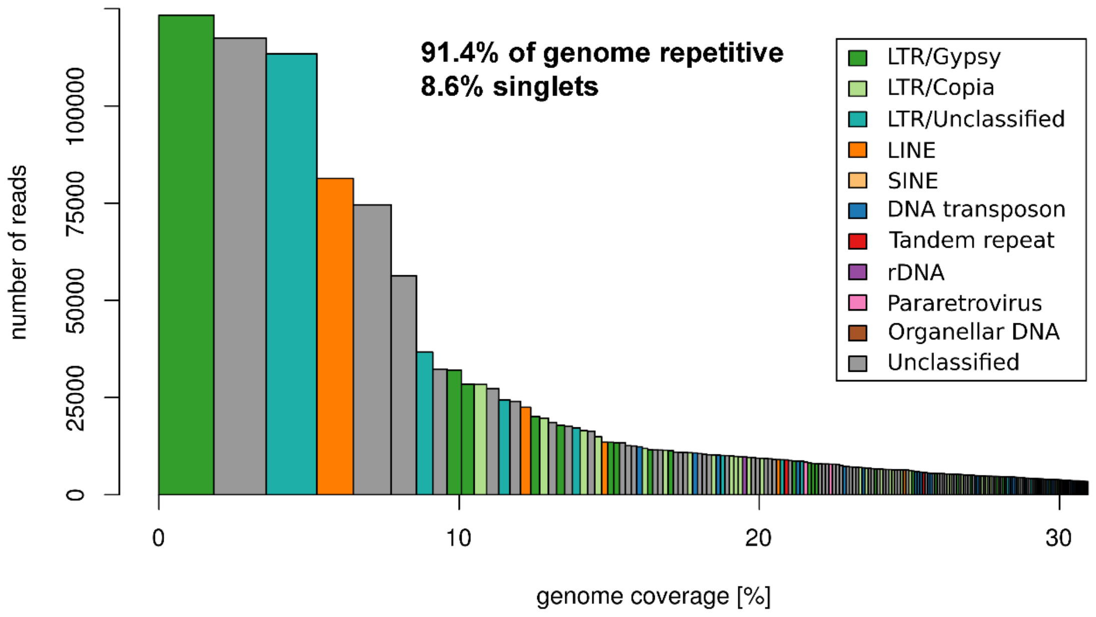


**Supplementary Figure 1**. RepeatExplorer output and repeat composition of superclusters according to RepeatExplorer analysis in *Vaccinium macrocarpon*. The graph is reproduced from data with permission from Sultana et al., 2017. (A) RepeatExplorer output without any treatment from RepeatExplorer analysis (B) Assignment of repeat types to the most abundant repeat clusters. Each column belongs to a specific supercluster and the corresponding color represents the types of repetitive DNA. Height of the column shows the number of reads and width represents the genome proportion for each supercluster.

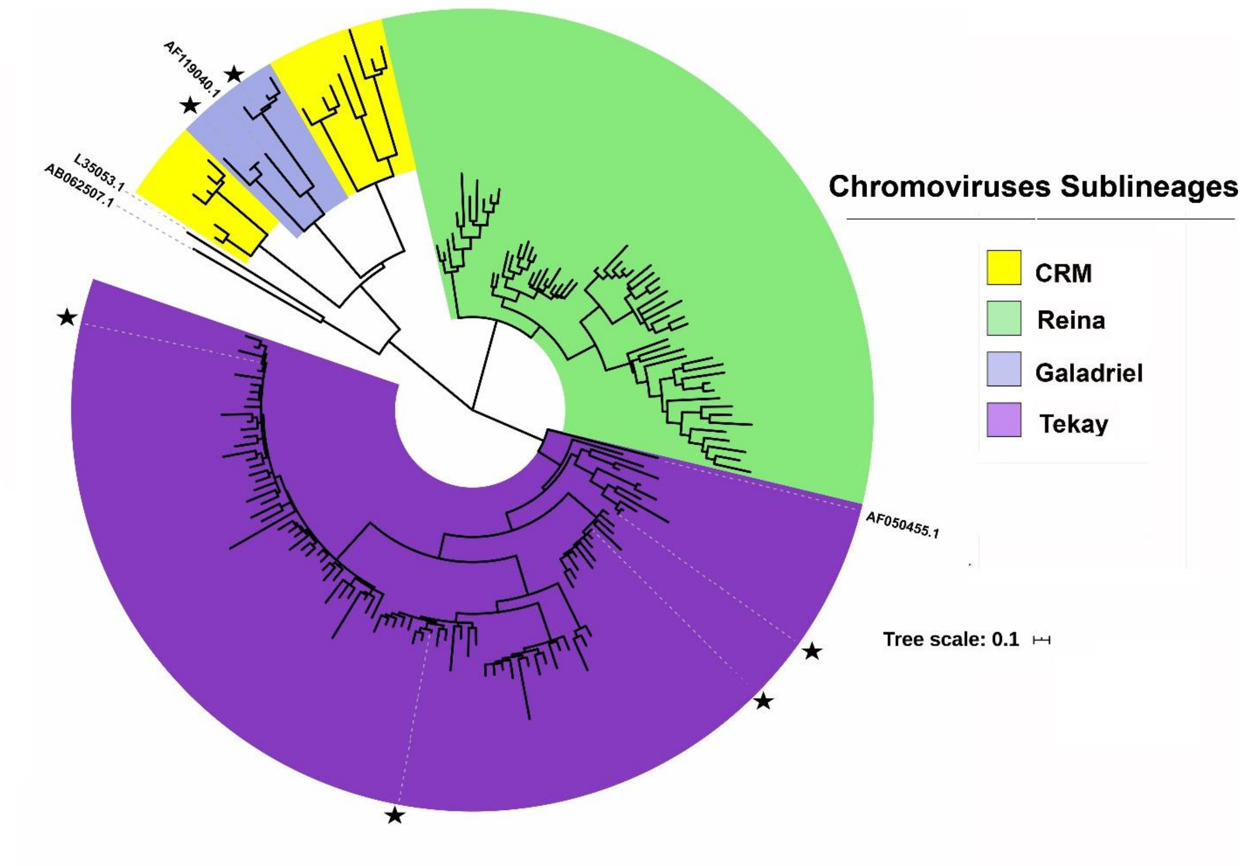


**Supplementary Figure 2**: Phylogenetic analysis of chromodomain (CHD) sequences identified from the genome assembly of *V. macrocarpon*. The tree was contructed using the Randomized Axelerated Maximum Likelihood RAxML method (Stamatakis et al., 2014). Scale bar of the tree is shown below. Highlighted branches (star) refer to CHD sequences of representative full-length sequences of chromoviruses reconstructed from *V. macrocarpon* Ilumina short reads. Outgroup (L35053 and B062507 from fungus genome *Magnaporthe grisea*) and reference chromodomain sequences (AF119040.1 for Galadriel and AF050455.1 for Tekay) were obtained from NCBI. Details of the outgroup sequences are recorded in Supplementary Table 3.

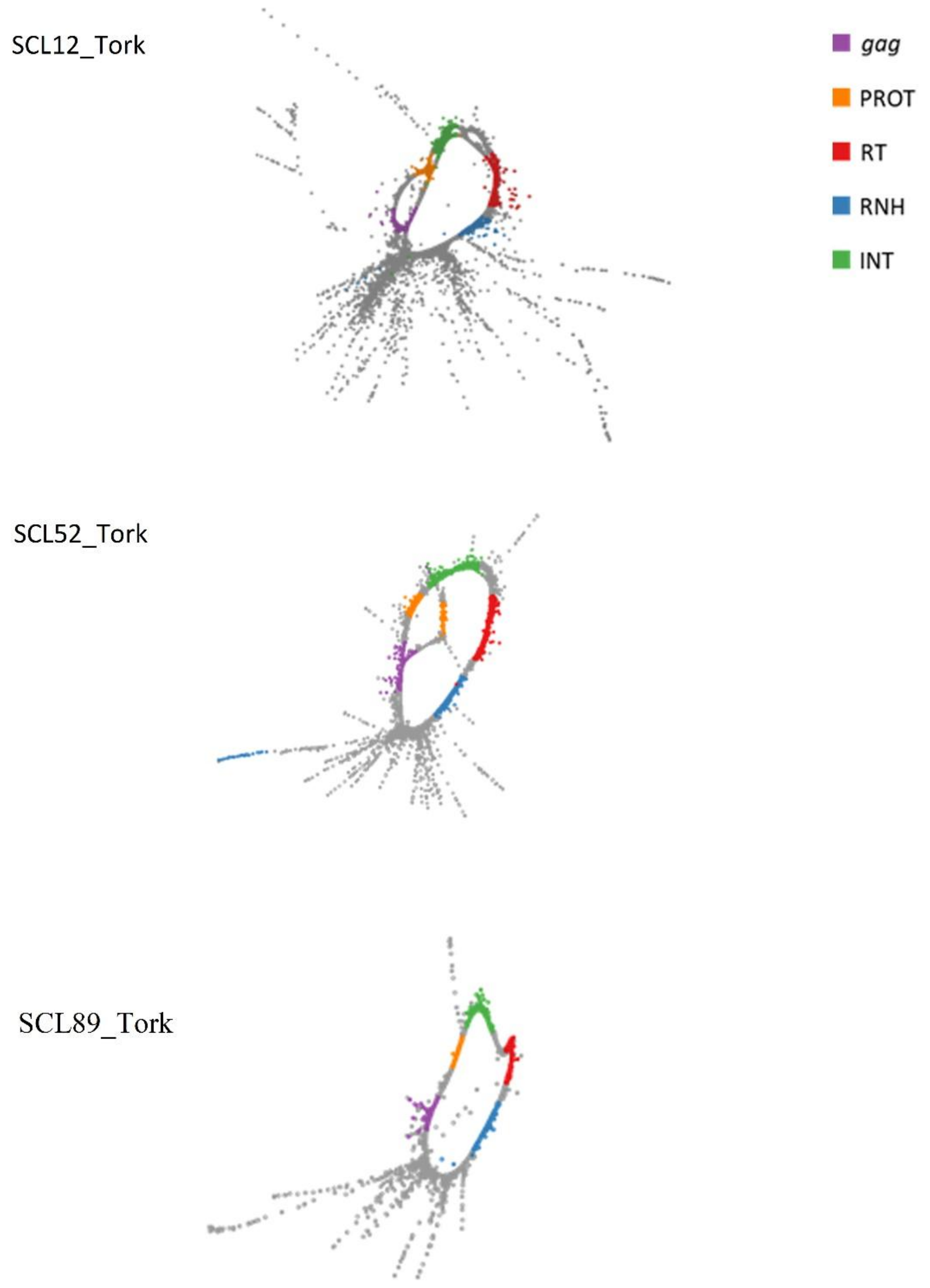


**Supplementary Figure 3.** Circular graph structure of selected Ty1-*copia* Tork elements with complete structural features. gag = *Gag* domain, PROT = protease, RT = reverse transcriptase domain, RNH = RNase H domain, , INT = integrase domain.

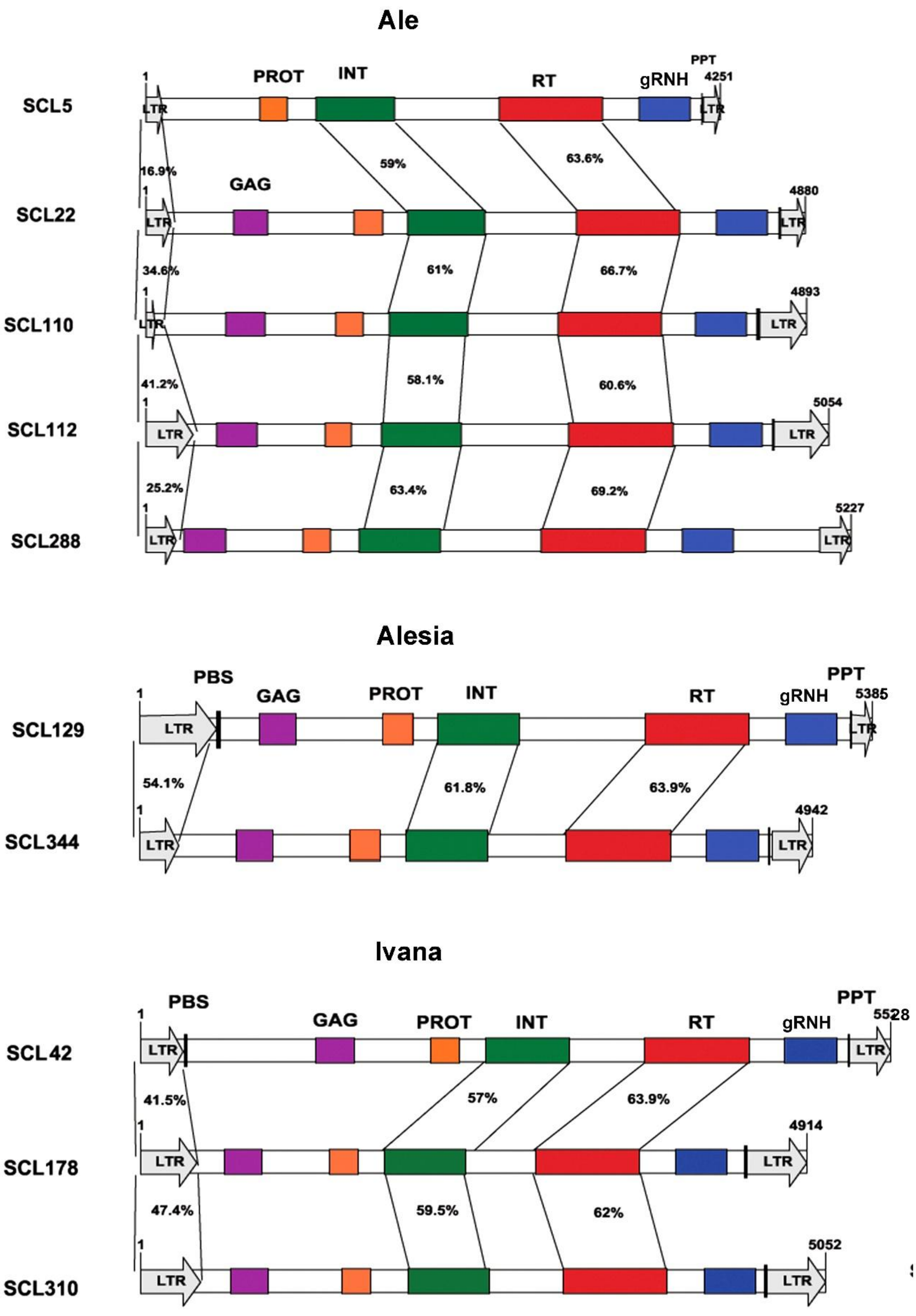

Ale
SCL5
SCL22
SCL110
SCL112
SCL288
PROT
INT
RT
gRNH
PPT
GAG
LTR
1
4251
4880
4893
5054
5227
16.9%
34.6%
41.2%
25.2%
59%
63.6%
61%
66.7%
58.1%
60.6%
63.4%
69.2%
Alesia
SCL129
SCL344
PBS
GAG
PROT
INT
RT
gRNH
PPT
LTR
1
5385
4942
54.1%
61.8%
63.9%
Ivana
SCL42
SCL178
SCL310
PBS
GAG
PROT
INT
RT
gRNH
PPT
LTR
1
5528
4914
5052
41.5%
47.4%
57%
63.9%
59.5%
62%

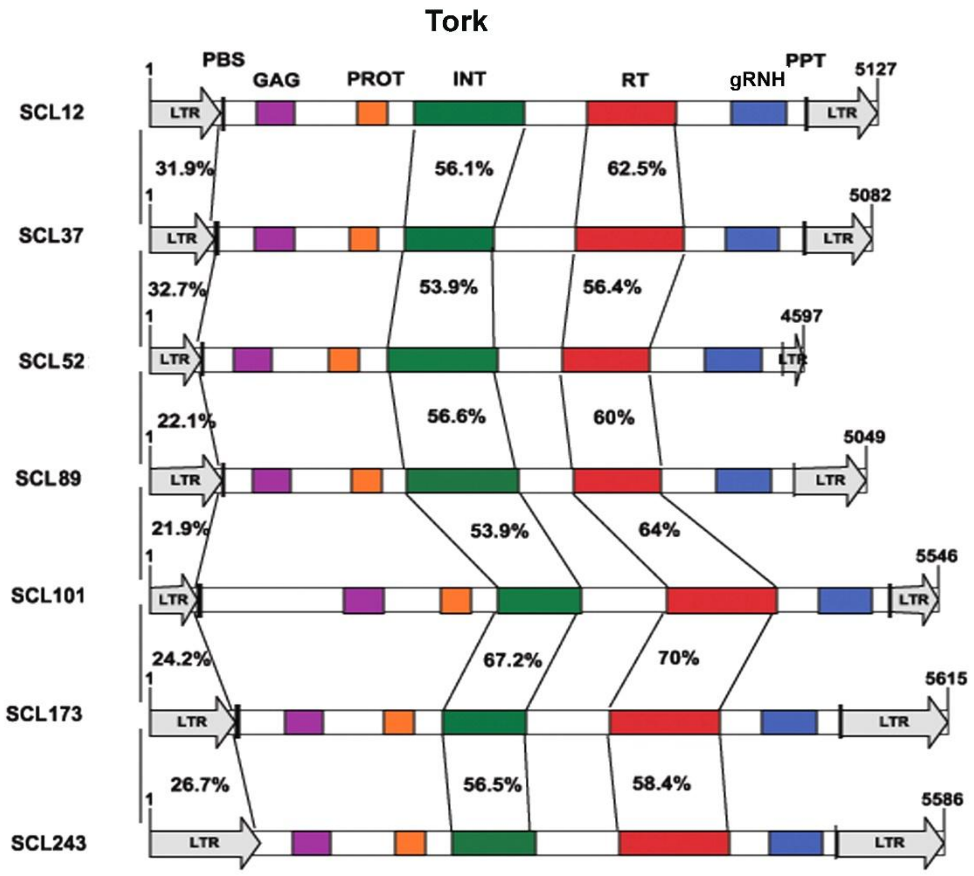
Tork
PBS
GAG
PROT
INT
RT
gRNH
PPT
SCL12
SCL37
SCL52
SCL89
SCL101
SCL173
SCL243
LTR
1
5127
5082
4597
5049
5546
5615
5586
31.9%
56.1%
62.5%
32.7%
53.9%
56.4%
22.1%
56.6%
60%
21.9%
53.9%
64%
24.2%
67.2%
70%
26.7%
56.5%
58.4%

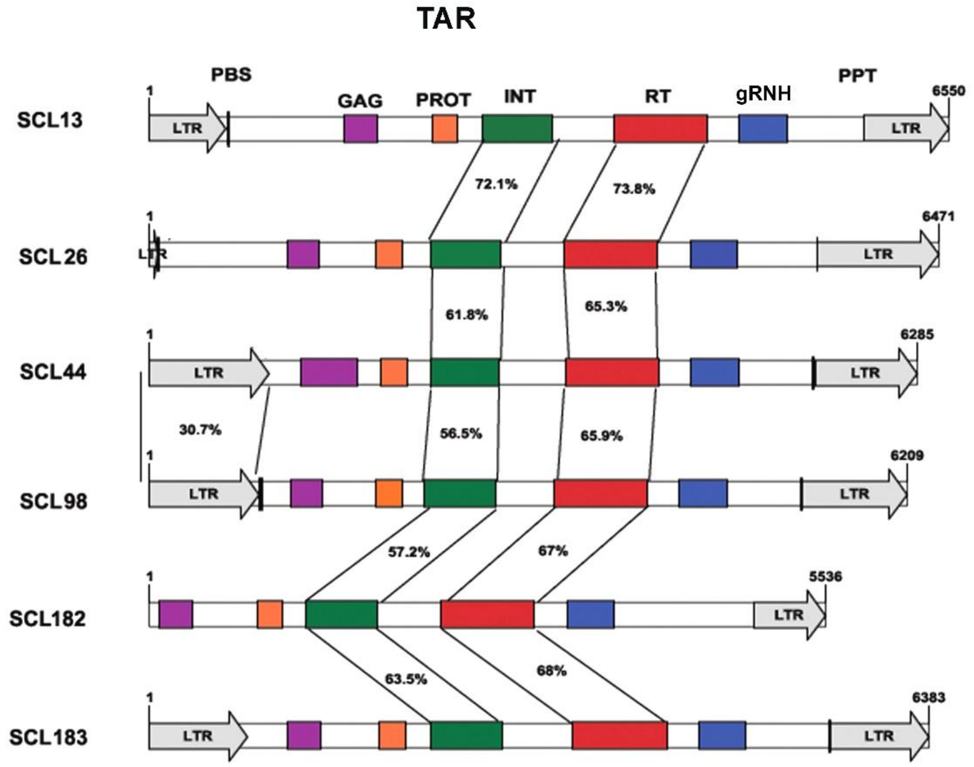
TAR
PBS
GAG
PROT
INT
RT
gRNH
PPT
SCL13
SCL26
SCL44
SCL98
SCL182
SCL183
LTR
1
6550
6471
6285
6209
5536
6383
72.1%
73.8%
61.8%
65.3%
30.7%
56.5%
65.9%
57.2%
67%
63.5%
68%

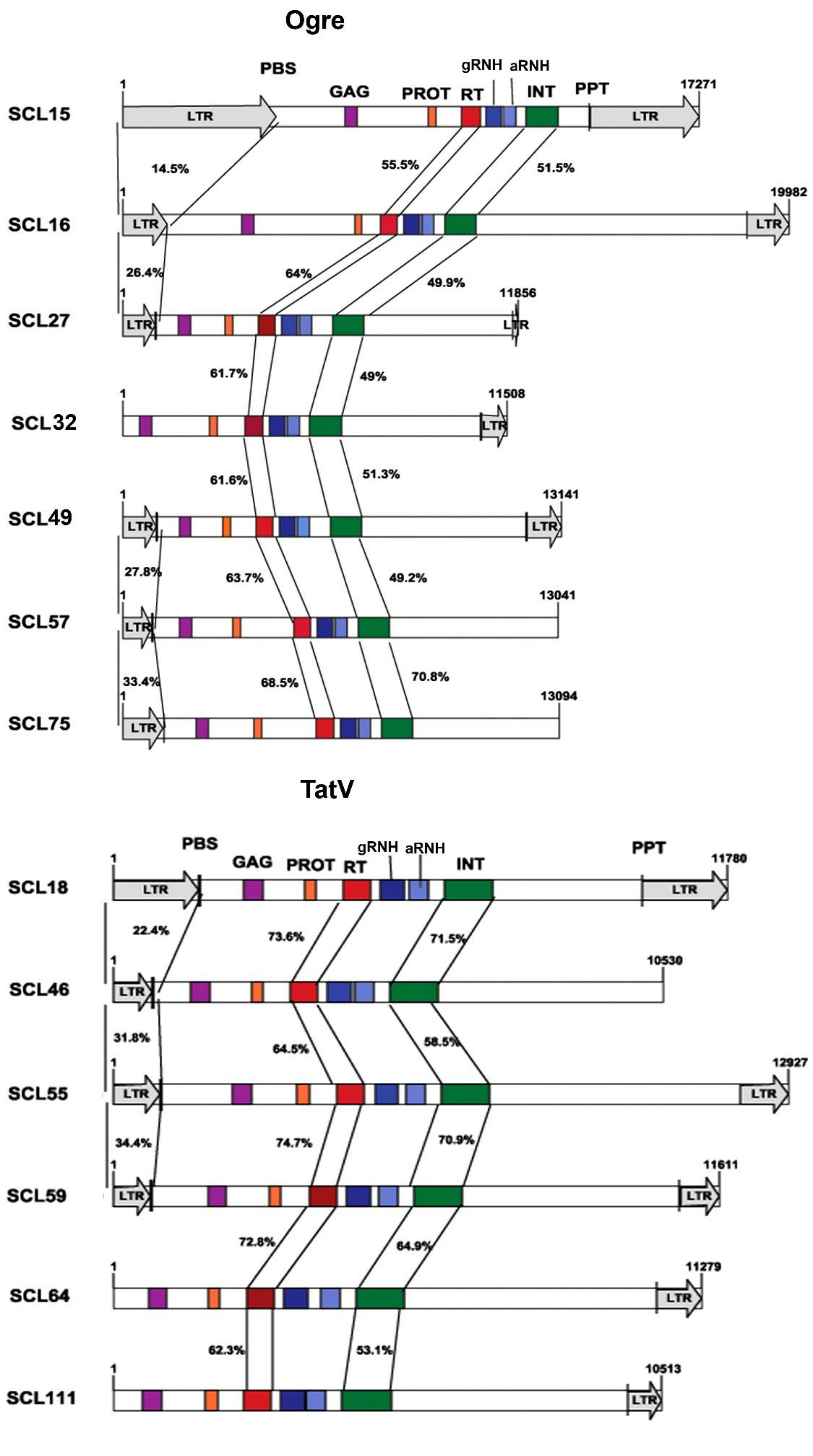
Ogre
PBS
GAG
PROT
RT
gRNH
aRNH
INT
PPT
LTR
SCL15
SCL16
SCL27
SCL32
SCL49
SCL57
SCL75
1
17271
19982
11856
11508
13141
13041
13094
14.5%
55.5%
51.5%
26.4%
64%
49.9%
61.7%
49%
61.6%
51.3%
27.8%
63.7%
49.2%
33.4%
68.5%
70.8%
TatV
SCL18
SCL46
SCL55
SCL59
SCL64
SCL111
11780
10530
12927
11611
11279
10513
22.4%
73.6%
71.5%
31.8%
64.5%
58.5%
34.4%
74.7%
70.9%
72.8%
64.9%
62.3%
53.1%

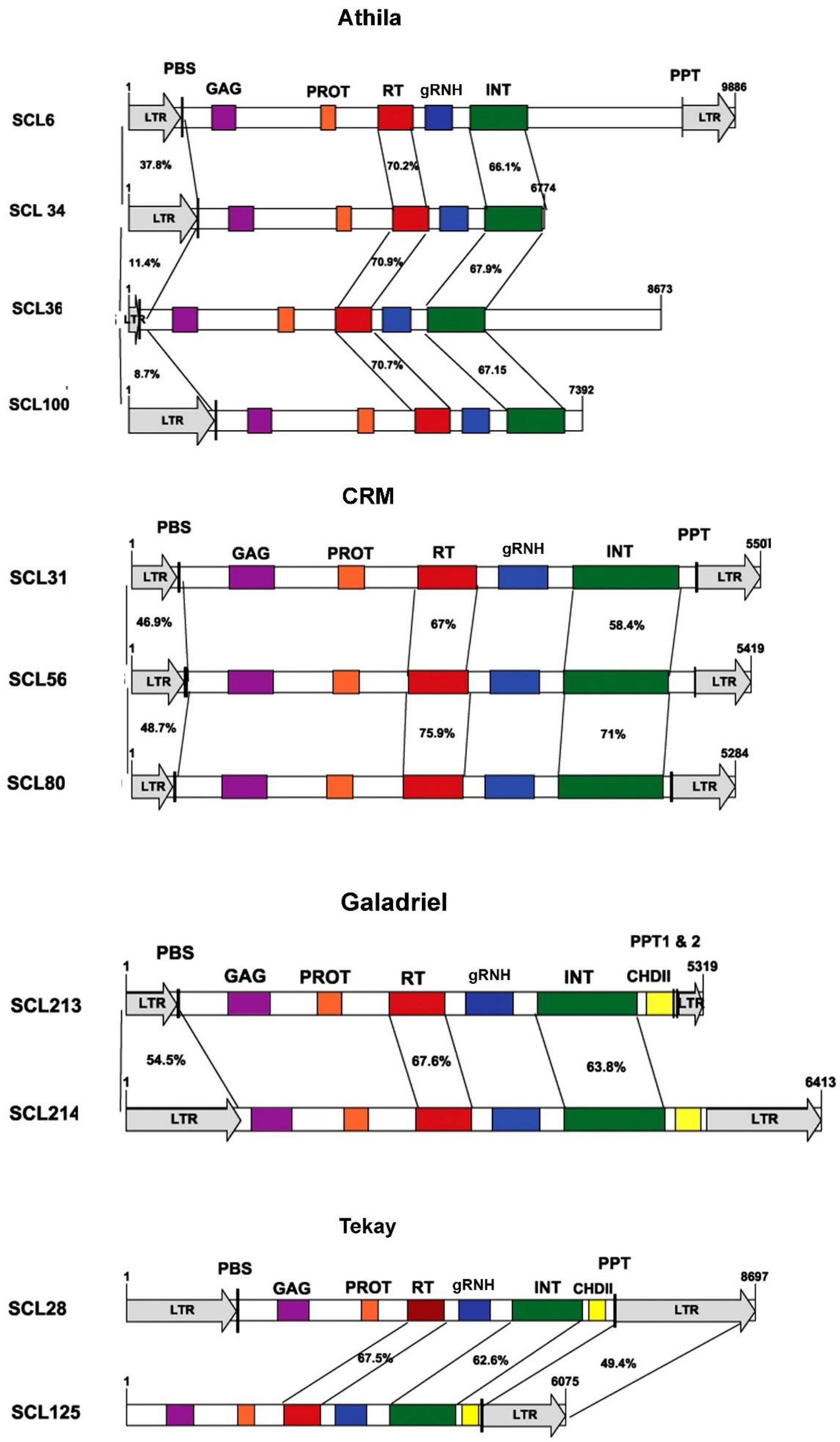


**Supplementary Figure 4.** Structural diversity of Ty1-*copia* and Ty3-*gypsy* retrotransposons in *V. macrocarpon*. Long terminal repeats (LTRs) are represented as gray colored open arrows (intact). Length of the arrow represents the length of the respective LTR. Please note the absence of the 3' LTR region in SCL182, SCL36, SCL46, SCL57, SCL75 and SCL100 and 5' LTR in SCL32, SCL64, SCL111, and SCL125. The black vertical lines adjacent to the LTRs represent the primer binding site (PBS) and polypurine tract (PPT). Structural features of *gag* and the four genes within *pol* are depicted as color-coded boxes: GAG = *gag*, PR = protease, RT = reverse transcriptase domain, gRNH = RNase H domain, aRNH = archaeal RNase H domain, INT = integrase domain, CHDII = chromodomain. Pairwise similarities for 5' LTR, RT and INT domain are provided among all the full-length elements.

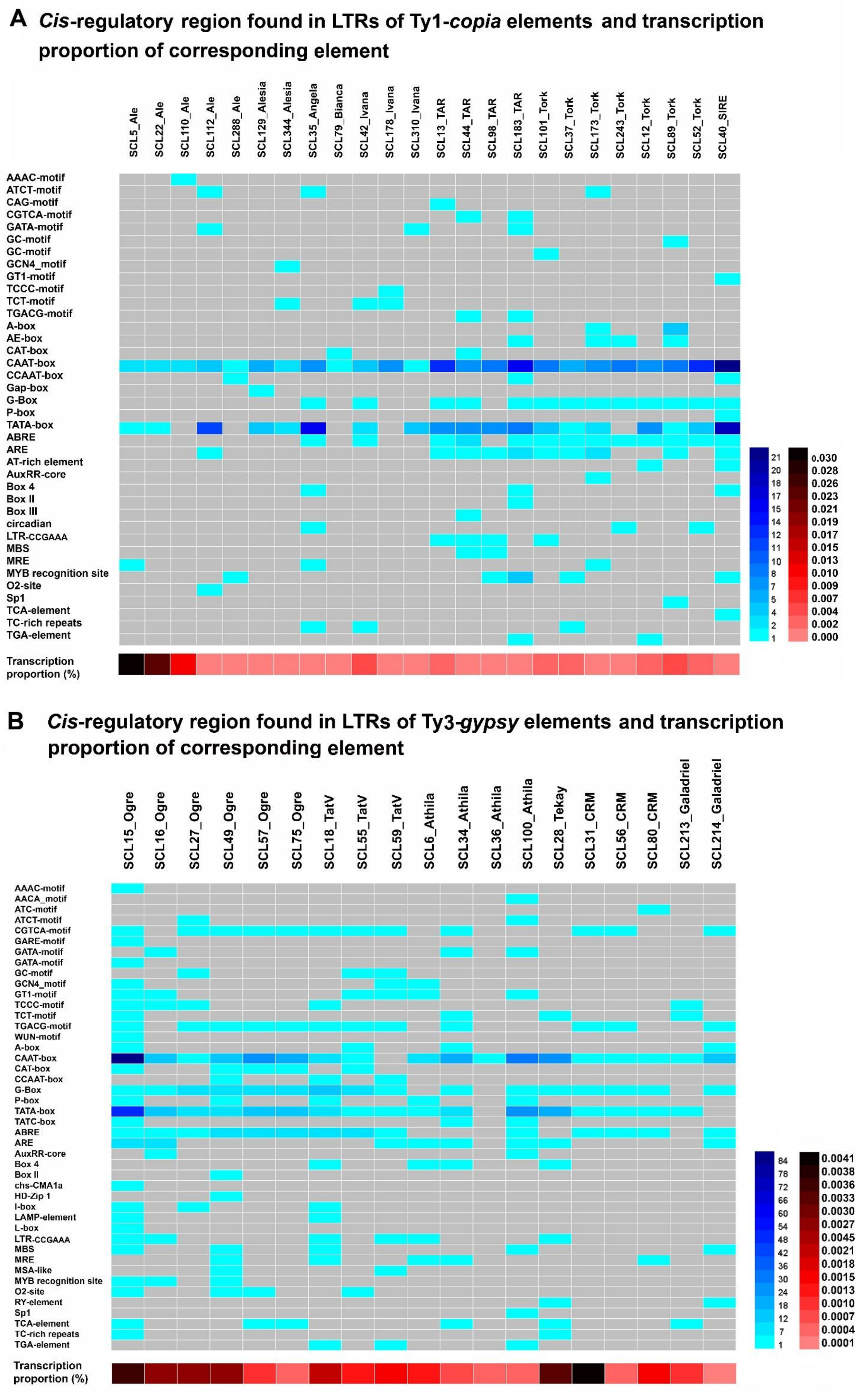


**Supplementary Figure 5.** *Cis*-regulatory elements in the LTRs of the *V. macrocarpon* full-length Ty1-*copia* and Ty3-*gypsy* LTR retrotransposons their corresponding transcriptome proportion. Heatmap representation of the number of *cis*-regulatory elements in the 24 representative (A) Ty1-*copia* and (B) the 19 Ty3-*gypsy* elements. Motifs were detected using The Plant Care database (http://bioinformatics.psb.ugent.be/webtools/plantcare/html). Color gradient in the scale bar on the right corresponds to the number of sequence motifs observed and transcriptome proportion observed. Details on motif abbreviations and potential function are presented in Supplementary Table 5.1 and 5.2.

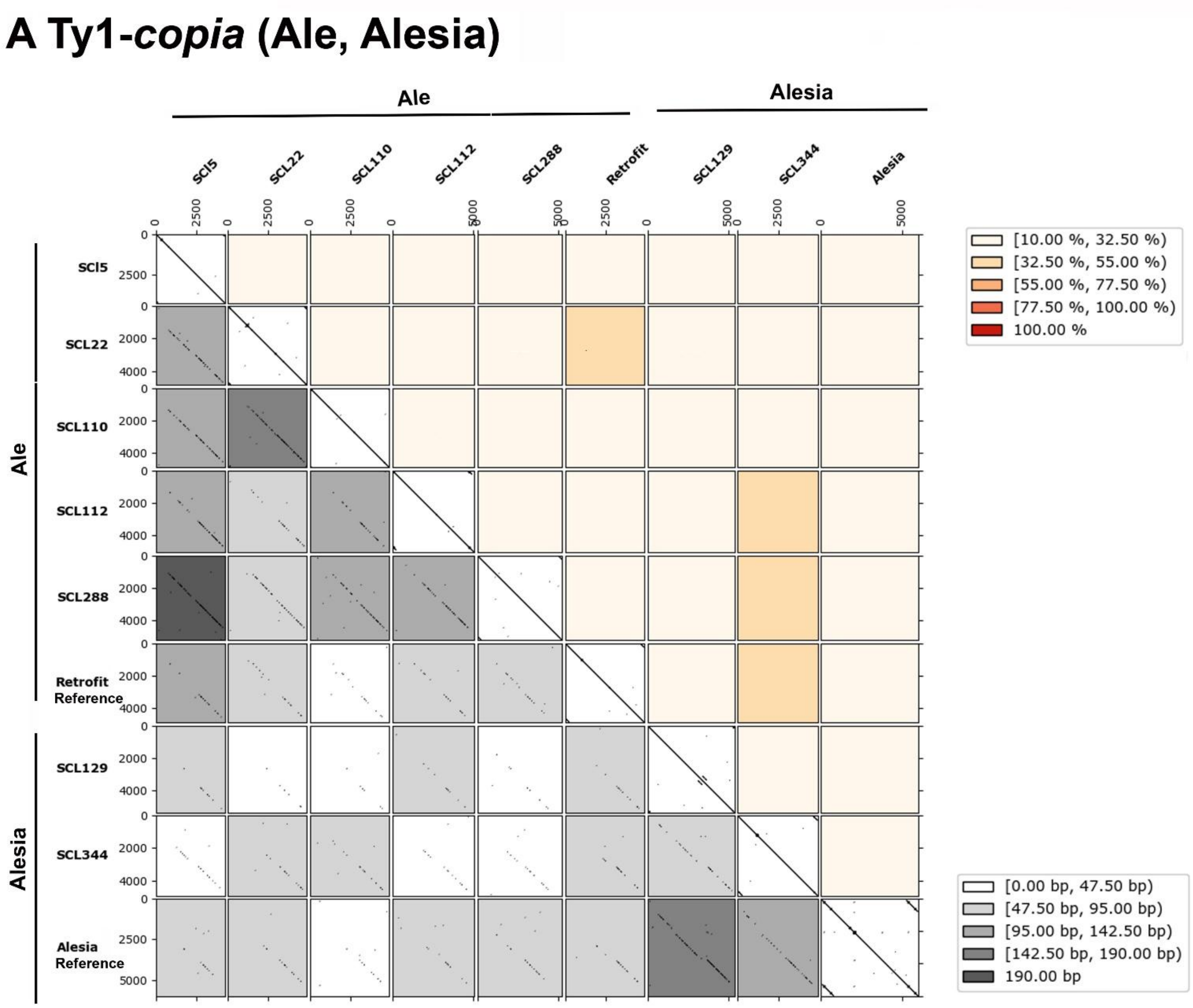
A Ty1-copia (Ale, Alesia)
Ale
Alesia
SCl5
SCL22
SCL110
SCL112
SCL288
Retrofit
SCL129
SCL344
Alesia
Retrofit Reference
Alesia Reference
[10.00 %, 32.50 %)
[32.50 %, 55.00 %)
[55.00 %, 77.50 %)
[77.50 %, 100.00 %)
100.00 %
[0.00 bp, 47.50 bp)
[47.50 bp, 95.00 bp)
[95.00 bp, 142.50 bp)
[142.50 bp, 190.00 bp)
190.00 bp

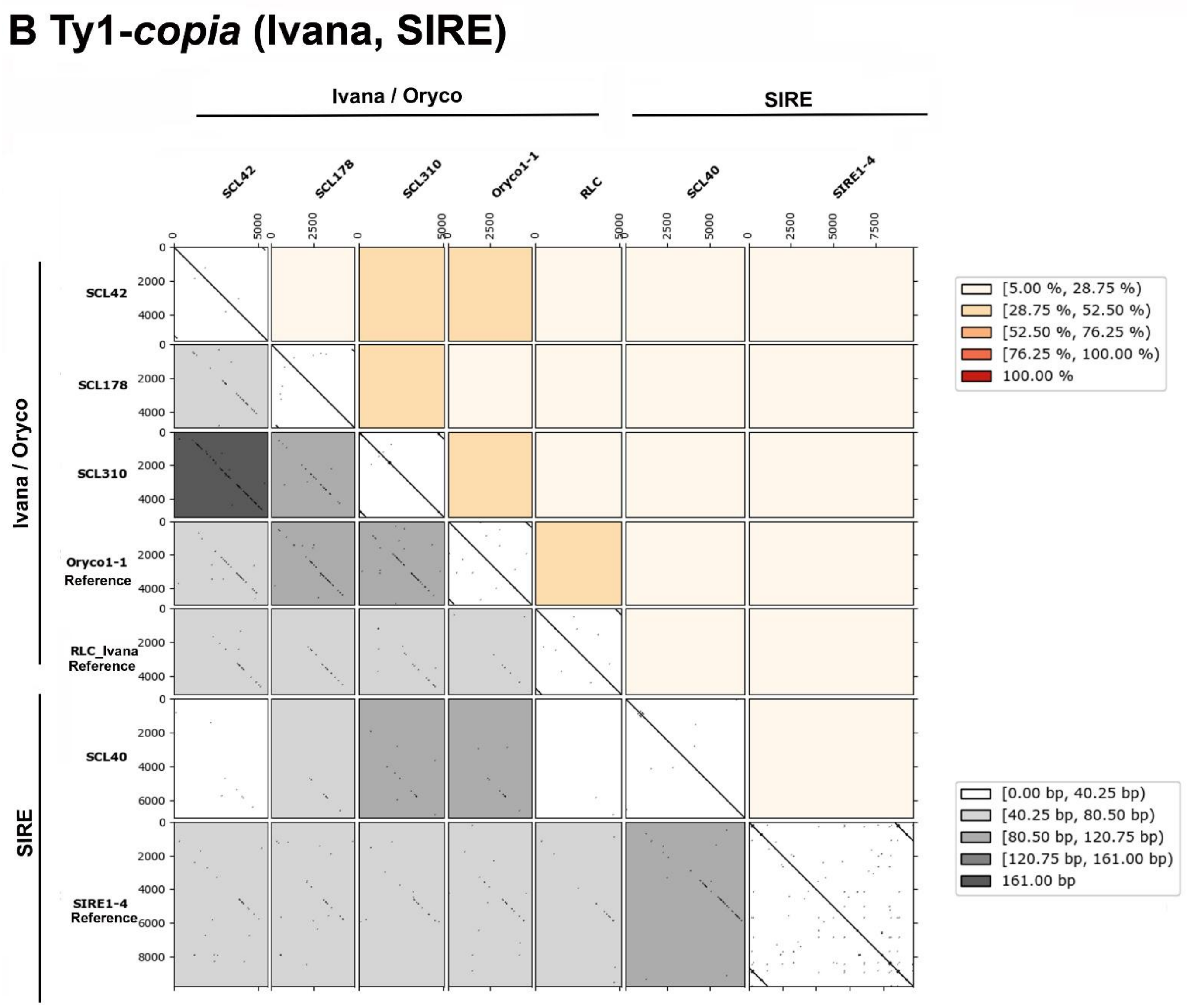
B Ty1-copia (Ivana, SIRE)
Ivana / Oryco
SIRE
SCL42
SCL178
SCL310
Oryco1-1
RLC
SCL40
SIRE1-4
Oryco1-1 Reference
RLC_Ivana Reference
SIRE1-4 Reference
[5.00 %, 28.75 %)
[28.75 %, 52.50 %)
[52.50 %, 76.25 %)
[76.25 %, 100.00 %)
100.00 %
[0.00 bp, 40.25 bp)
[40.25 bp, 80.50 bp)
[80.50 bp, 120.75 bp)
[120.75 bp, 161.00 bp)
161.00 bp

#### C *Ty1-copia* (Tork, TAR, Angela, Bianca)

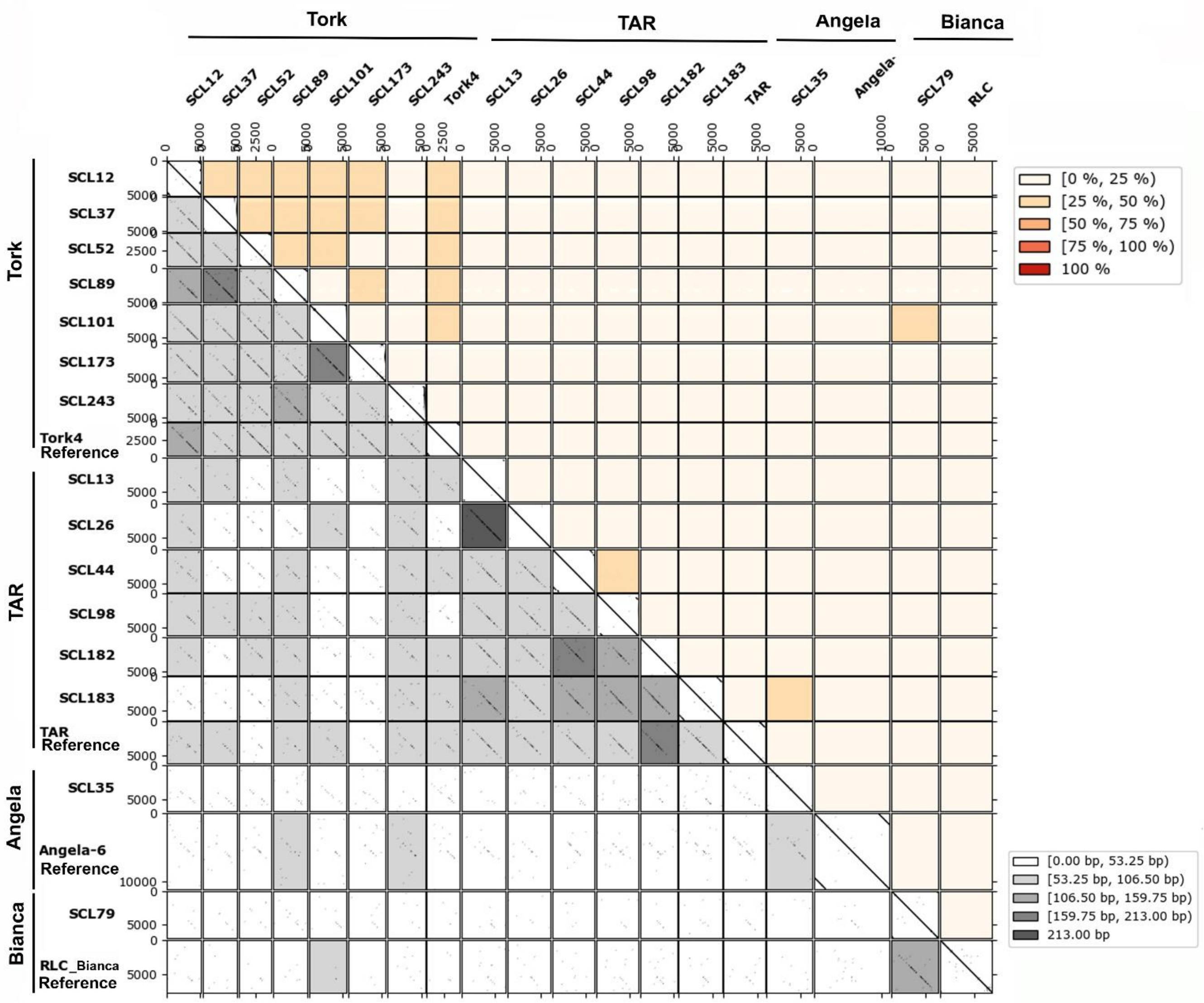

### D Ty3-*gypsy* (Non-chromovirus)

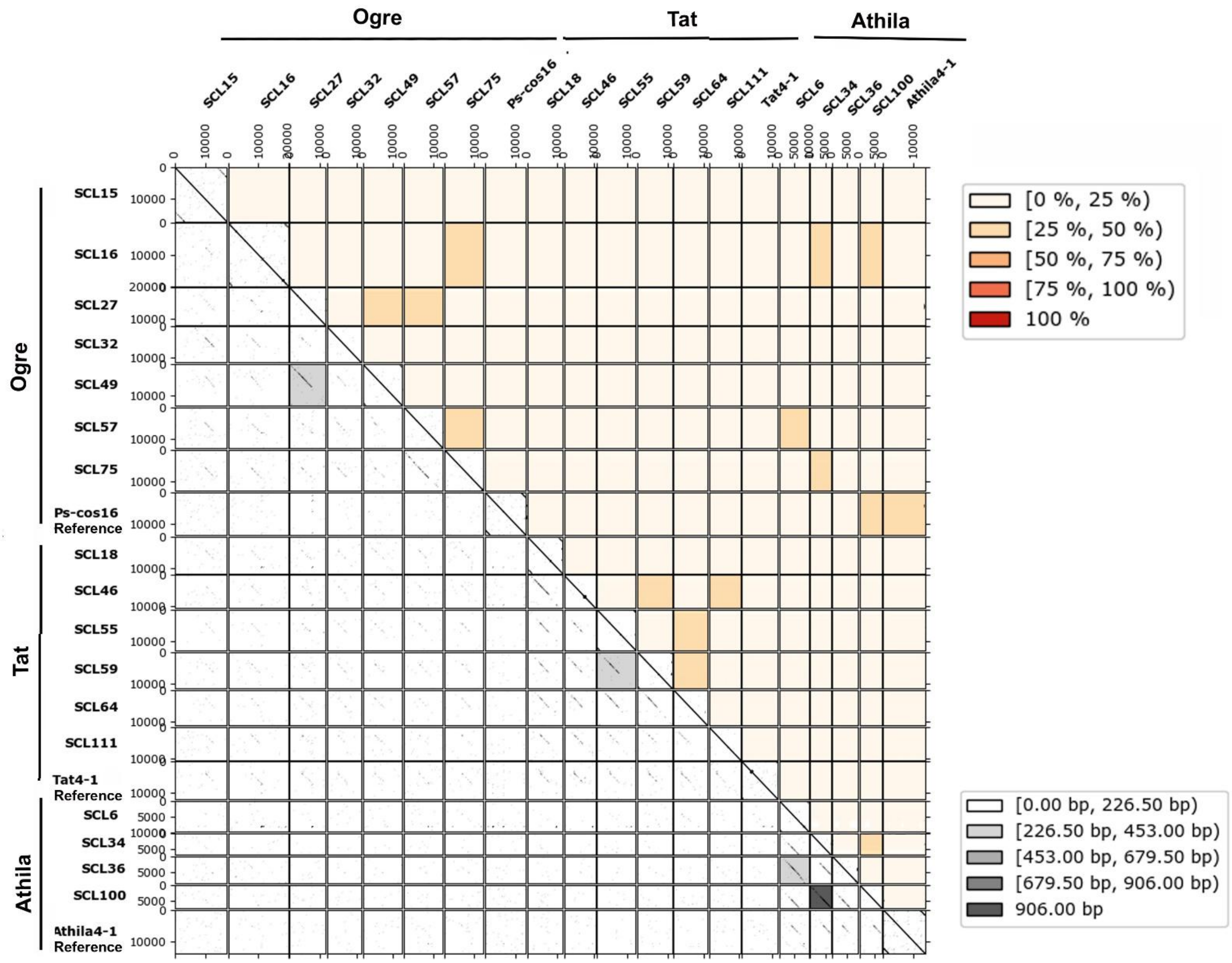

### E Ty3-*gypsy* (Chromovirus)

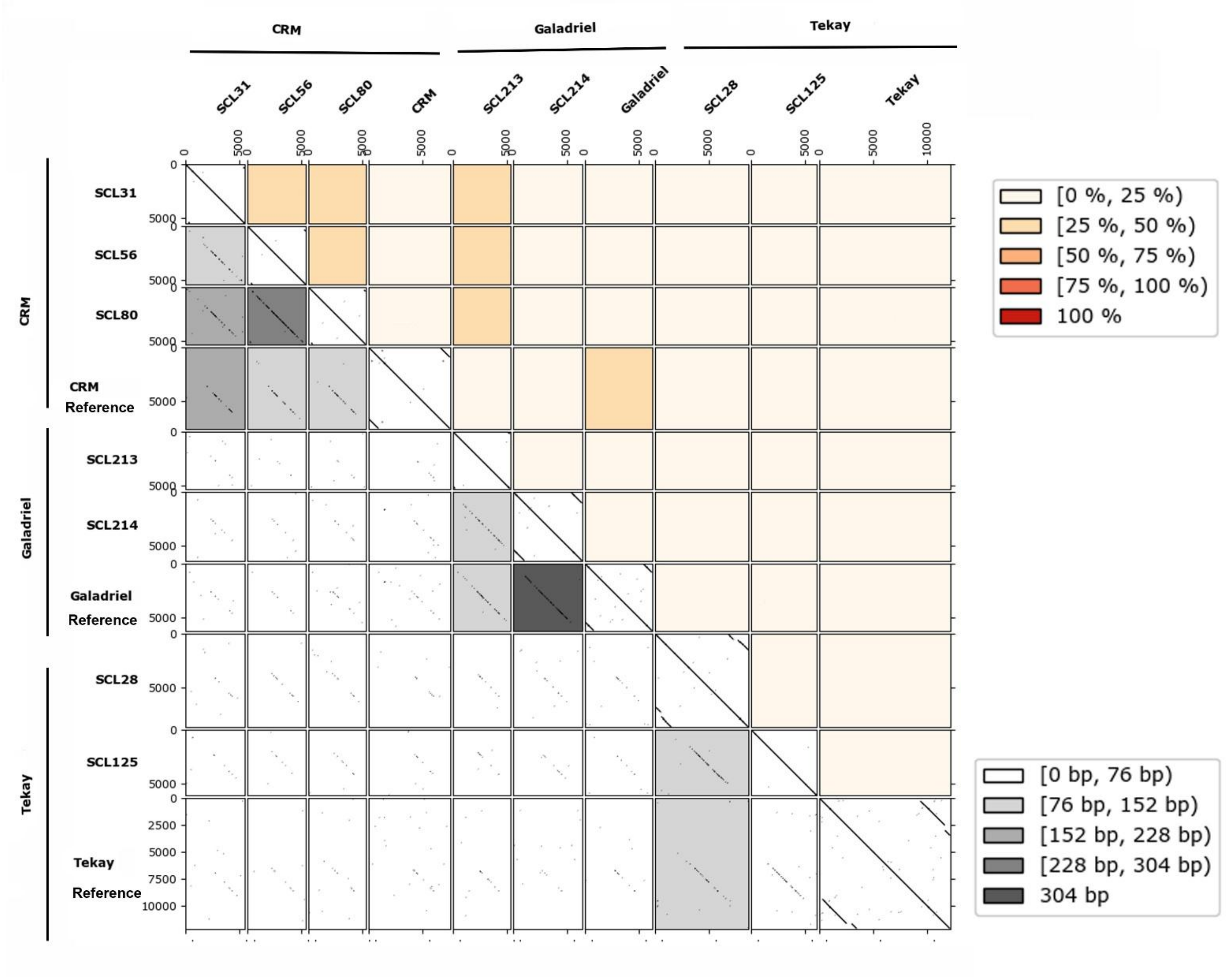


**Supplementary Figure 6.** All against all dotplot comparison of phylogenetically closely related full-length elements of the lineages of the Ty1-*copia* and the Ty3-*gypsy* superfamily in the *Vaccinium macrocarpon* genome. For each lineage of (A-C) the Ty1-*copia* and (D-E) the Ty3-*gypsy* superfamily, dotplots were calculated for the phylogenetically closely related reconstructed full-length elements including reference elements from Gypsy Database 2.0 (Llorens et al. 2011), TREP Database (http://botserv2.uzh.ch/kelldata/trep-db/index.html) and NCBI. Detailed information of these reference elements were provided in Supplementary Table 3. Dotplots were calculated with a wordsize of 25 (-k 25) and with allowing 6 mismatches (-S 6) using FlexiDot (Seibt et al. 2019). In the lower half, dotplots are presented. Each rectangle represents a pairwise sequence comparison, with the main diagonal containing self-comparisons. In these self dotplots, LTRs are visible as short diagonals in the upper right and the lower left corner. Dotplots are shaded according to the longest uninterrupted match (= longest diagonal). In the upper half, pairwise identity of the 5' LTRs are given and shaded accordingly.

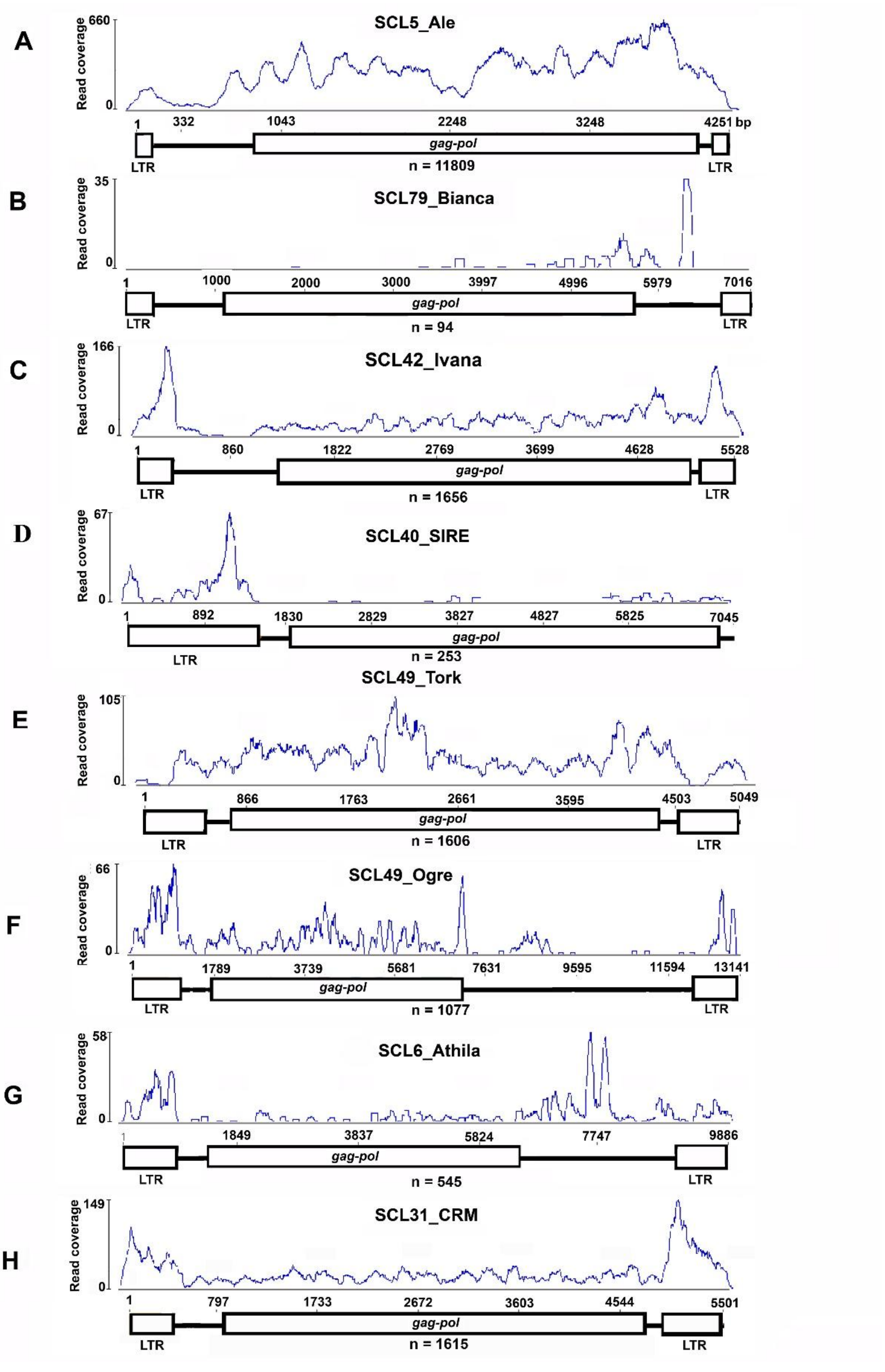


**Supplementary Figure 7.** Transcriptome read mapping to selected elements representing the different lineages of the Ty1-*copia* and Ty3-*gypsy* superfamilies in *V. macrocarpon* as evidence for transcriptional activity. Transcript read coverage for representative *in silico* full-length elements of (A-E) Ty1-*copia* and (F-H) Ty3-*gypsy* retrotransposons are shown with a schematic element structure underneath. Read coverage axis is presented in linear scale (n = total transcript count).
