## Supplementary Tables 1-8 for "Genome-wide analysis of long terminal repeat retrotransposons from the cranberry *Vaccinium macrocarpon*": Supplementary Table 3_Total number of reverse transcriptase (RT) protein sequences.docx

Supplementary Table 3: Total number of reverse transcriptase (RT) protein sequences identified using the RepeatExplorer protein domain tools on the assembled genome of *V. macrocarpon*.

| Classification according to Repeatexplorer (class/superfamily/clade) | Family/Lineage | Total number of RT protein sequences | Percentage of total RT sequences (%) | Number of clustered sequences^1^ |
| --- | --- | --- | --- | --- |
| Class_I\|LTR\|Ty1\|copia\| |  |  |  |  |
|  | Ale | 587 | 27.32 | 109 |
|  | Alesia | 8 | 0.37 | 1 |
|  | Angela | 7 | 0.33 | 7 |
|  | Bianca | 34 | 1.58 | 3 |
|  | Ikeros | 27 | 1.26 | 3 |
|  | Ivana | 86 | 4.00 | 12 |
|  | SIRE | 21 | 0.98 | 7 |
|  | TAR | 105 | 4.89 | 22 |
|  | Tork | 142 | 6.61 | 12 |
| Class_I\|LTR\|Ty3\|gypsy\| |  |  |  |  |
|  | Chromovirus\|CRM | 105 | 4.89 | 34 |
|  | Chromovirus\|Galadriel | 13 | 0.60 | 4 |
|  | Chromovirus\|Reina | 25 | 1.16 | 15 |
|  | chromovirus\|Tekay | 116 | 5.40 | 28 |
|  | non-chromovirus\|OTA\|Athila | 134 | 6.24 | 25 |
|  | non-chromovirus\|OTA\|Ogre/Tat | 739 | 34.39 | 183 |

^1^at 90% similarity threshold after removing ambiguous sequences
