## Supplementary Tables 1-8 for "Genome-wide analysis of long terminal repeat retrotransposons from the cranberry *Vaccinium macrocarpon*": Supplementary Table 4_Total number of chromodomain (CHD) protein sequences.docx

Supplementary Table 4: Total number of chromodomain (CHD) protein sequences identified in the assembled genome of *V. macrocarpon* with the RepeatExplorer tool DANTE.

| Chromovirus clade | Total number of chromodomain (CHD) sequences | Percentage of total CHD sequences (%) | Number of clustered sequences^1^ |
| --- | --- | --- | --- |
| CRM | 21 | 5.69 | 20 |
| Galadriel | 18 | 4.88 | 5 |
| Reina | 133 | 36.04 | 61 |
| Tekay | 197 | 53.39 | 92 |

^1^at 90% similarity threshold after removing ambiguous sequences
