## Supplementary Data File 1 for "Genome-wide analysis of long terminal repeat retrotransposons from the cranberry *Vaccinium macrocarpon*": Read me_Information.docx

**Information about this Full-length LTR-sequences**

**Title: Genome-wide analysis of Ty3-*gypsy* and Ty1-*copia* long terminal repeat (LTR) retrotransposons from *Vaccinium macrocarpon* Aiton**

Nusrat Sultana^1^, SedatSerçe^2^, Gerhard Menzel^3^, Tony Heitkam^3^, Thomas Schmidt^3^

^1^ Department of Botany, Faculty of Life and Earth Sciences, Jagannath University, Dhaka 1100, Bangladesh

^2^ Department of Agricultural Genetic Engineering, Ayhan Şahenk Faculty of Agricultural Sciences and Technologies, Niğde Ömer Halisdemir University, 51240, Niğde, Turkey.

^3^ Institute of Botany, Technische Universität Dresden, D-01062 Dresden, Germany

Submitted in **Biology** (ISSN 2079-7737; CODEN: BBSIBX), an international peer-reviewed open access journal of biological sciences published quarterly online by MDPI.

**Table 1.** Repeat specific name of full-length reconstructed sequences of Ty3-*gypsy* and Ty1/*Copia* elements from *V. macrocarpon* cultivar ‘Ben Lear’

| Elements name | Elements |
| --- | --- |
| SCL5_Ale |  |
| SCL22_Ale |  |
| SCL110_Ale |  |
| SCL112_Ale |  |
| SCL288_Ale |  |
| SCL129_Alesia |  |
| SCL344_Alesia |  |
| SCL35_Angela |  |
| SCL79_Bianca |  |
| SCL42_Ivana |  |
| SCL178_Ivana |  |
| SCL310_Ivana |  |
| SCL13_TAR |  |
| SCL26_TAR |  |
| SCL44_TAR |  |
| SCL98_TAR |  |
| SCL182_TAR | Ty1-*Copia* |
| SCL183_TAR |  |
| SCL101_Tork |  |
| SCL37_Tork |  |
| SCL173_Tork |  |
| SCL243_Tork |  |
| SCL12_Tork |  |
| SCL89_Tork |  |
| SCL52_Tork |  |
| SCL15_Ogre |  |
| SCL16_Ogre |  |
| SCL27_Ogre |  |
| SCL32_Ogre |  |
| SCL49_Ogre |  |
| SCL57_Ogre |  |
| SCL75_Ogre |  |
| SCL18_TatV |  |
| SCL46_TatV |  |
| SCL55_TatV | Ty3-*gypsy* |
| SCL59_TatV |  |
| SCL64_TatV |  |
| SCL111_TatV |  |
| SCL6_Athila |  |
| SCL34_Athila |  |
| SCL36_Athila |  |
| SCL100_Athila |  |
| SCL125_Tekay |  |
| SCL28_Tekay |  |
| SCL31_CRM |  |
| SCL56_CRM |  |
| SCL80_CRM |  |
| SCL213_Galadriel |  |
| SCL214_Galadriel |  |
